# Graph-guided assembly for novel HLA allele discovery

**DOI:** 10.1101/138826

**Authors:** Heewook Lee, Carl Kingsford

**Affiliations:** Computational Biology Department, School of Computer Science, Carnegie Mellon University

## Abstract

Accurate typing of human leukocyte antigen (HLA), a histocompatibility test, is important because HLA genes play various roles in immune responses, and have also been shown to be associated with many diseases such as cancer. The current gold standard for HLA typing uses DNA sequencing technology combined with sequence enrichment techniques using specially designed primers or probes, causing it to be slow and labor-intensive. Although there exist enrichment-free computational methods that use various types of sequencing data, hyper-polymorphism found in HLA region of the human genome makes it challenging to type HLA genes with high accuracy from whole genome sequencing data. Furthermore, these methods are database-matching approaches where their output is inherently limited by the completeness of already known types, forcing them to find the best matching known alleles from a database, thereby causing them to be unsuitable for discovery of rare or novel alleles. In order to ensure both high accuracy as well as the ability to type novel alleles, we have developed a graph-guided assembly technique for classical HLA genes, which is capable of assembling phased, full-length haplotype sequences of typing exons given high-coverage (>30-fold) whole genome sequencing data. Our method delivers highly accurate HLA typing, comparable to the current state-of-the-art database-matching methods. We also demonstrate that our method can type novel alleles by experimenting on various data including simulated, Illumina Platinum Genomes, and 1000 Genomes data.

## Introduction

Human leukocyte antigen (HLA) genes are crucial to the regulation of immune system as they encode for the major histocompatibility complex (MHC) consisting of cell surface proteins that control the adaptive immune response. HLA genes are also known to play important roles in transplant rejection, autoimmune disorders [1] and cancer [2, 3]. For these reasons, accurate HLA typing is important both in clinical and research settings. HLA typing is considered challenging because of the hyper-polymorphic nature of the HLA region in human genome. Such high polymorphism in the HLA region is thought to be maintained by strong balancing selection promoting genetic diversity [4, 5]. Especially with personal genome sequencing becoming widely common, better computational methods are needed to provide rapid and inexpensive typing with high accuracy.

Traditionally, HLA typing or categorization was done by more laborious serology-based methods to screen for HLA antibodies in a donor/receiver pair. With the birth of DNA sequencing and polymerase chain reaction (PCR), molecular typing assays such as specific oligonucleotide probe hybridization (SSOP), sequencespecific primer amplification (SSP), and sequence-based typing (SBT) have been developed [6]. The SBT method can be either used with Sanger sequencing or NGS techniques. By using specific primers to perform target enrichment prior to sequencing, SBT delivers accurate and reliable typing of HLA alleles. However, all of the above molecular typing assays require a specially designed set of probes/primers, and they are labor intensive, low throughput, and costly.

With an increasing availability of personal WGS services, an availability of accurate computational HLA typing methods that do not require additional experiments can be valuable. Challenges in computational HLA typing are mainly driven by the high level of polymorphism found in the HLA region in the human genome. There are over 30 genes that are maintained in the IPD-IMGT/HLA database [7] and there are 6-8 classical HLA genes (HLA-A, -B, -C, -DQA, -DQB, -DRB) routinely used for HLA typing in clinical settings. More than 15,000 known alleles (just for these classical genes) have been reported in the database and the number of alleles is growing rapidly (Figure 1). Also, the known alleles share high sequence similarities, where many alleles just differ by a base-pair substitution. Thus, it is challenging to correctly pinpoint an individual’s HLA types among the known alleles using WGS data [8].

**Figure 1.**
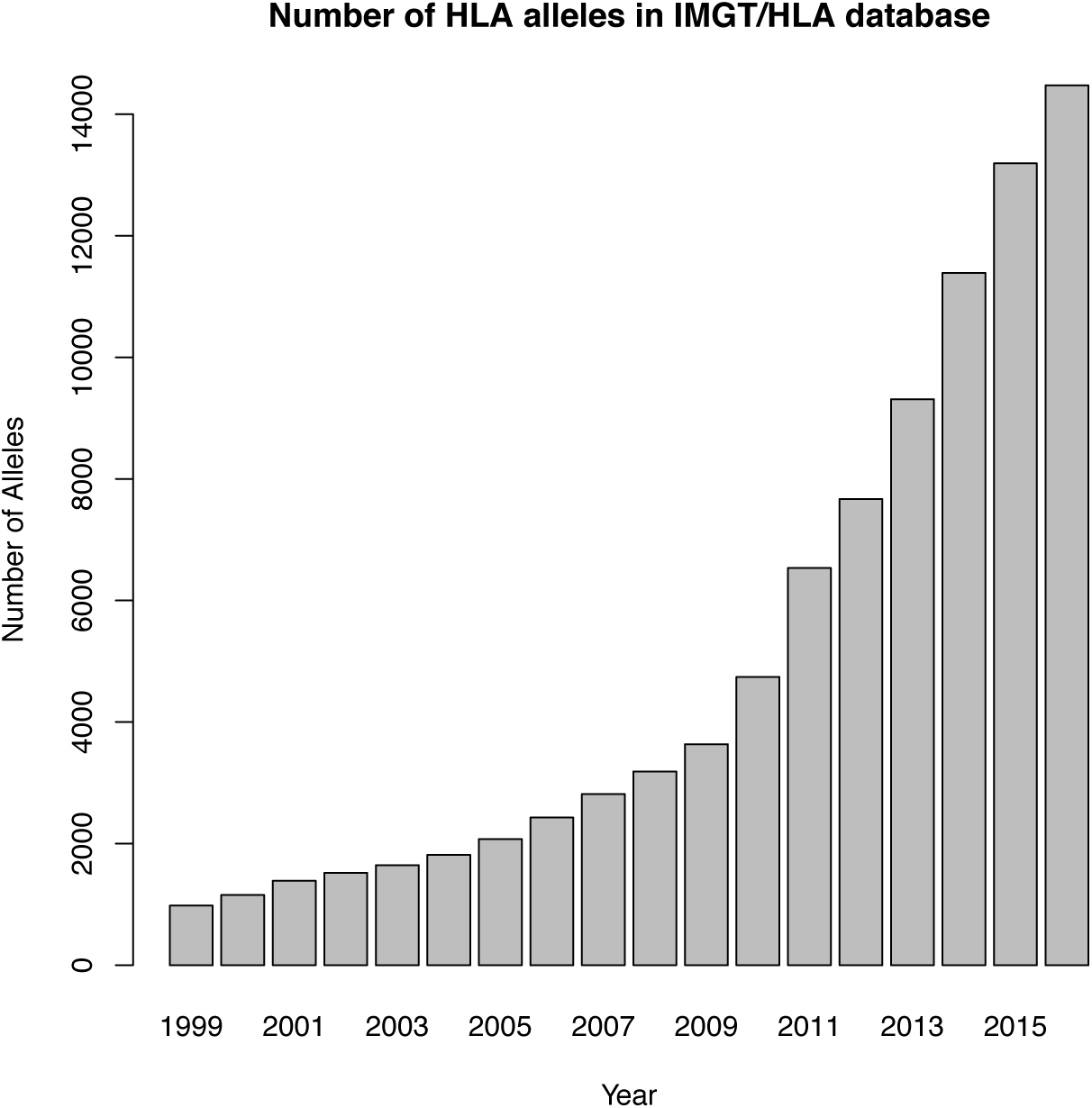
The number of alleles in the IPD-IMGT/HLA Database by year from 1999 to 2016. The database releases updates 4 times a year (January, April, July, and October) and the plot is based on number of alleles from all the April releases reported on the statistics page of the IPD-IMGT/HLA website (http://www.ebi.ac.uk/ipd/imgt/hla/stats.html).

Previously developed enrichment-free computational methods can use whole genome sequencing (WGS), whole exome sequencing (WES), or transcriptome sequencing (RNA-seq) without the use of HLA-enriched data as opposed to the SBT. However, they do not provide typing accuracy comparable to what the SBT provides [9].

These methods either use one or both of two major techniques—alignment and assembly—to accurately compare reads to correct HLA genes and infer allele types. Alignment-based methods try to correctly assign NGS reads to HLA loci then infer typing alleles using widely used probabilistic genotyping techniques. Assembly-based methods first construct longer-than-read contigs using *de novo* assembly techniques and search for best matches among known alleles and direct assembly of haplotypes by traditionally available *de novo* or reference-based assemblers are severely confounded by the high level of polymorphisms. `PHLAT` [10] uses probabilistic SNP calling techniques and also models phasing by looking at reads covering neighboring variant sites. `HLA-VBSeq` [11] uses variational bayesian inference to correctly assign reads to alleles based on read-to-allele alignment and outputs a sorted list of alleles by the number of reads assigned. `HLA*PRG` [12] performs an alignment of extracted reads likely coming from HLA region to population reference graphs [13] that encode all known alleles and outputs the most likely alleles from the database.

One common ground for the enrichment-free computational HLA typing methods is that they are all driven by the *finding-the-nearest-match* paradigm; their goal is to find the best matching alleles to HLA genes of a test individual in a preexisting database of known entries. Given sequencing data of an individual, such a typing scheme outputs the best matching alleles for each HLA gene. This typing strategy is limited by the completeness of the collection of known alleles as it cannot detect novel alleles missing in the database of known types. We collectively refer to these approaches as *database-matching* methods. Novel alleles can possibly have protein coding changes which may have more profound impact in the context of organ transplant and disease association. One might argue that there are already many known alleles and that the chance of finding novel alleles is low. However, the number of known alleles in the IPD-IMGT/HLA database is still increasing rapidly (Figure 1). There is continuing effort among immunogenetics communities to study rare and novel alleles. For example, the International HLA and Immunogenetics Workshop has been organizing projects to investigate and collect rare and novel alleles since their 15th workshop in 2002. Immunogenetics-related journals such as International Journal of Immunogenetics, and HLA (formerly known as Tissue Antigens) both have a dedicated section where new alleles are announced in every issue.

For these reasons, it is important to be able to recover HLA sequences at 1-bp resolution to enable novel allele discovery as similarly done in SBT. In order to achieve this goal, we present a graph-guided assembly technique called `Kourami` that constructs full sequences for the peptide binding domain (exons 2 and 3 for class I and exon 2 for class II HLA genes–regions typed by the SBT methods) by using a modified partial ordered graph (POG) [14] as a guide. Our method is the first method that directly assemble both haplotypes of HLA genes rather than to infer the best matching alleles in the database. For known alleles, we show that Kourami can correctly type with high accuracy (>98%), equalling that of the state-of-the-art database matching method, across various WGS datasets such as simulated data, Illumina Platinum Genomes, and high coverage WGS from 1000 Genomes project. At the same time, `Kourami` only takes a fraction of time compared to other available methods with a moderate use of memory.

In addition, `Kourami` is the first HLA typer to be able to assemble novel alleles that do not appear in the database. It does this by treating the HLA typing problem as an instance of graph-guided assembly, where the known alleles are combined into a graph that is used to guide the assembly of new alleles. `Kourami` therefore also represents an early example of how a population of reference sequences can be used during genome assembly. We systematically show that Kourami is very accurate in its ability to construct novel alleles by performing leave-one-out experiments where a known allele is artificially removed from the allele database. `Kourami` is able to reconstruct 98% of these alleles perfectly.

## Materials and Methods

### HLA typing nomenclature

The current HLA allele nomenclature [15] uses a hierarchical numbering system with 4 major levels of hierarchies. From the highest to the lowest category, it annotates allele groups (2-digit resolution), protein sequence (4-digit resolution), exon sequence (6-digit resolution), and intron sequence (8-digit resolution). For example, if two alleles encode an identical protein, they will have the same numbers for the first 2 levels (4-digit) of hierarchies. In practice, HLA typing is often carried out at either the protein or exon level. Furthermore, the current gold standard, SBT, types just the exons that are responsible for encoding the peptide binding domain (exons 2 and 3 for class I genes and exon 2 for class II genes). Using only the subset of exons creates ambiguous alleles where two or more alleles share identical sequence over these exons but differ in other regions. These ambiguous sequences are grouped as a 6-digit ‘G’ allele. Similarly, 4-digit ‘P’ grouping is used for the alleles that share same amino acid sequence over these exons. Our method provides fully assembled sequence covering these exons and also outputs 6-digit ‘G’ resolution typing result by selecting known alleles that have the smallest edit distance to the assembled sequences. Similar to many other HLA tools, we focus on the routinely typed classical genes (HLA-A, -B, -C, -DQA1,-DQB1, -DRB1).

### Overview of method

Our method takes an advantage of partial order graphs to capture all known alleles and further modifies the graph to include variants found in the sequencing data for graph to include paths of true alleles. An overview of our method is illustrated in Figure 2, and the major steps are labeled from (a) to (e). More details are given in “Materials and Methods”. We first create a comprehensive reference panel from a combined multiple sequence alignment (MSA) of both full-length and exon-only known alleles for each HLA locus (step a). Reads mapped to all known HLA loci in the human genome reference (GRCh38) are extracted (step b) and aligned to the comprehensive reference panel (step c). Gene-wise partial-ordered graphs are constructed using the combined MSAs and the alignments of the extracted reads are projected onto the graphs so that each read alignment is stored as a path in the graphs and read depths on edges naturally become edge weights (step d). When these read- or read-pair-backed paths connect 2 or more neighboring heterozygous sites of 2 alleles, they provide phasing information. During the alignment projection, the graphs are modified by adding nodes and edges to incorporate differences found by alignment such as substitutions and indels. Note that a sequence of an allele is encoded as a path through the entire graph. Finally, with the weighted graphs with alignment paths, we formulate the problem of constructing the best pair of HLA allele sequences as finding the pair of paths through the graph. When finding the pair, we consider consistent phasing information from the reads and coverage with a use of base quality scores. Additionally, the pair of paths may be identical to permit homozygous alleles.

**Figure 2.**
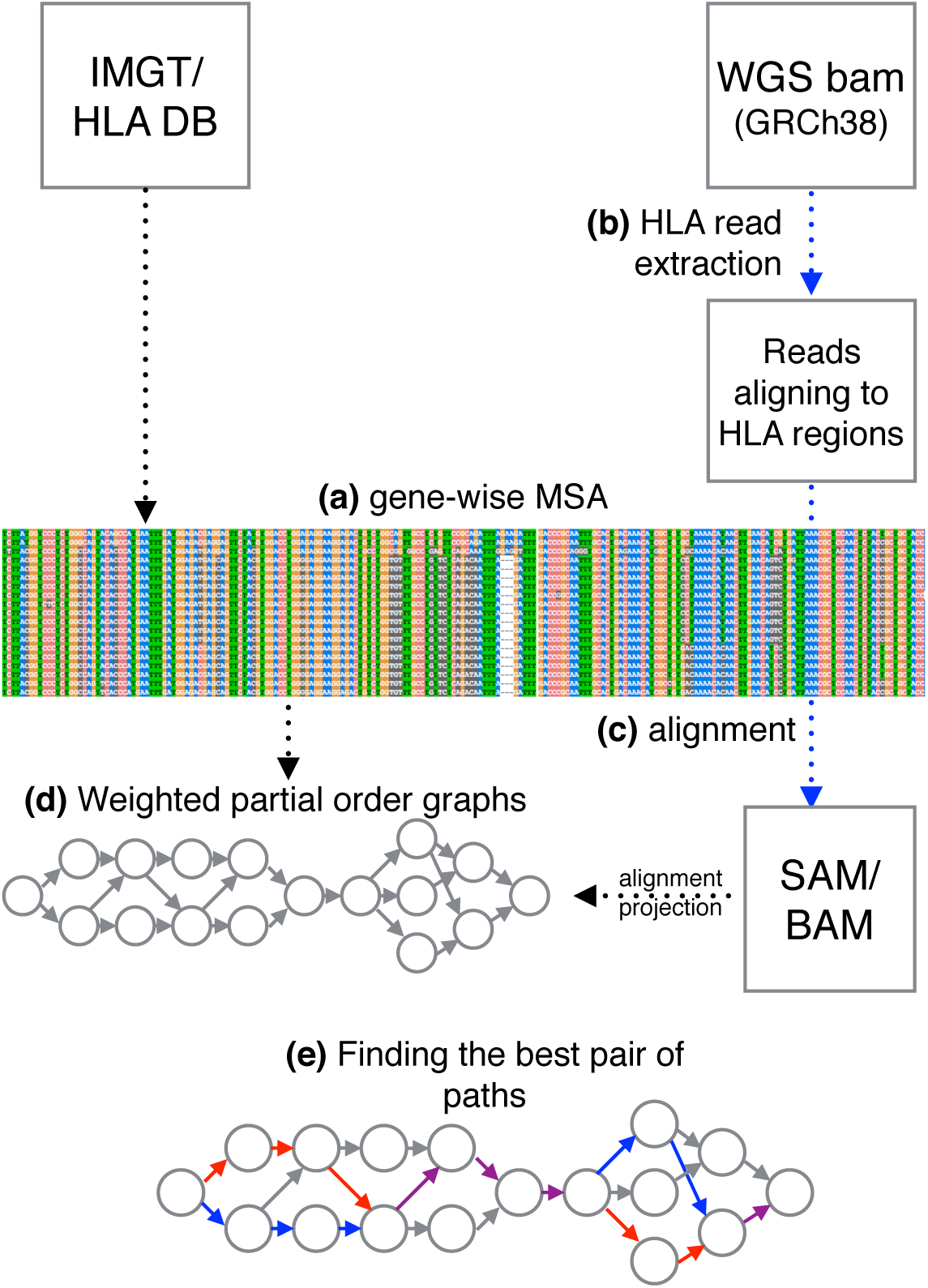
Overview of our method. (a) A gene-wise MSA is obtained from the IMGT/HLA database. The reads aligning to HLA regions are extracted (b) from the input bam and they are realigned (c) to the sequences in the MSA. (d) A POG is constructed from MSA and further modified via alignment projection. (e) Haplotype assembly of two alleles is obtained by finding two paths (drawn in red and blue;overlap in purple) through the graph.

### Input alignment and extraction of HLA reads

`Kourami` takes alignment of WGS to the human genome as an input in the BAM format. For many experiments used here, we used pre-computed alignments downloaded from Eurepean Bioinformatics Institute and Google Cloud Platform. In case of missing alignment files, we follow the 1000 Genomes procedures (see the GRCh38DH alignment readme file available from the 1000 Genome FTP server) to align reads using `BWA-kit` v0.7.15 [16] and further process the bam files using other tools such as `BioBamBam` [17] and `GATK` [18].

From the alignments, we extract paired-end reads aligned to all known HLA loci in chromosome 6, alternate sequences (ALT) of extended MHC (xMHC) regions, and HLA sequences (the complete set of coordinates used is in Table S1 in Supplementary Materials) included in the Human reference genome (hs38DH packaged in `BWA-kit` v0.7.15). In the GRCh38 assembly, regions that exhibit sufficient variability are represented in the primary chromosomal sequence as well as the ALT loci scaffolds.

### Known HLA alleles and construction of a comprehensive reference panel

The Immuno Polymorphism Database (IPD) maintains a periodically updated database of known HLA alleles in the IPD-IMGT/HLA database [7]. IPD-IMGT/HLA Release 3.24.0 (April, 2016) was used for all experiments here. A detailed breakdown of numbers of alleles included in this release is shown on Table 1. The other methods compared here use earlier versions of the database because the content of database is built into their software and there is no way to update or swap database at the user level. Using a later version of the database does not give advantages as long as the earlier version also contains the true alleles of testing individual.

**Table 1:** Number of known HLA alleles used (Release 3.24.0)

| Genes |  | Full-length | Total<br>(exonic + full-length) |
| --- | --- | --- | --- |
| Class I | A | 218 | 3399 |
|  | B | 337 | 4242 |
|  | C | 301 | 2950 |
| Class II | DQA1 | 45 | 69 |
|  | DQB1 | 27 | 911 |
|  | DRB1 | 40 | 1883 |

Full-length denotes the total of number full-length alleles in the release and total number includes the full-length alleles and the alleles with only exon sequences reported.

Many alleles in the database only contain partial sequences, often just covering few exons responsible for peptide binding domain of HLA genes (Table 1). For this reason, the IPD provides a set of pre-computed multiple sequence alignments (MSA) of full length alleles (*M_gene_*) and just the coding regions (*M_coding_*) separately for each HLA gene. Similarly to `HLA*PRG` [12], for each HLA gene, we combine these two MSAs by aligning them at corresponding columns in order to obtain a comprehensive reference panel of known alleles. This can help better recruit reads that span intron-exon junctions. The combined MSA (*M_panel_*) has the same number of rows as *M_coding_*. The number of columns in *M_panel_* is equal to the sum of the number of columns in *M_coding_* and the number of intronic columns in *M_gene_*. For each row in M_*coding*_, if the allele for the row has a corresponding row in *M_gene_*, intronic columns are inserted into *M_coding_*, otherwise, intronic columns of the reference allele in *M_gene_* are inserted. In addition to the HLA genes that are included in the IPD-IMGT/HLA database, non-polymorphic HLA genes DQA2 and DQB2, paralogous copies of DQA1 and DQB1 and often regarded as poorly polymorphic, are added to the reference panel as decoys to filter out reads originating from them aligning incorrectly to other class II genes. In our analysis, we noticed that reads coming from DQA2 or DQB2 can make the assembly of typing exons of class II genes difficult as previously reported [10].

### HLA-graph construction

In order to capture all information contained in *M_panel_* in a minimal manner as well as to allow flexibility to enable novel sequence discovery, we use partial order graphs, a compact graphical representation for MSA [14]. From each *M_panel_*, we can directly construct a gene-specific partial order graph similar to ones typically used in multiple sequence alignment [14, 19]. An example of a MSA of 3 known sequences (*M_panel_*) is shown in Figure 3(a). Each sequence is first drawn as a chain of vertices connected by directed edges (Figure 3(b)), where each vertex *v*_*i*_ represents a base symbol *b*_*v*_*i*__ (*b*_*v*_*i*__ ∈ {*A*, *C*, *G*, *T*, *N*, –}) and is positioned at column *i* in the graph. For each column, vertices with an identical base symbol at a column are merged as a single vertex and duplicate edges are removed (Figure 3(C),(d)). The gap symbol (‘-’) is used to restrict edges to connect vertices only from consecutive columns in the input MSA. An edge between two vertices (*e_v_i_, v_i+1__*) exists if *M_panel_* has a row with consecutive bases *b*_*v*_*i*__ and *b*_*v*_*i+1*__ at columns *i* and *i* + 1. It is important to note that this graph contains at least the same number of paths as the number of rows in *M_panel_* used to construct the graph. The graph often encodes a larger number of paths and such flexibility of the graph is the foundation which allows us to model this family of seqeunces and capture novel alleles. For example, a simple graph shown in Figure 3(d) encodes all sequences in the given MSA as well as AGGT-A, ACGTCA, and ACCTCA. Each path through the constructed graph encodes a possible allele.

**Figure 3.**
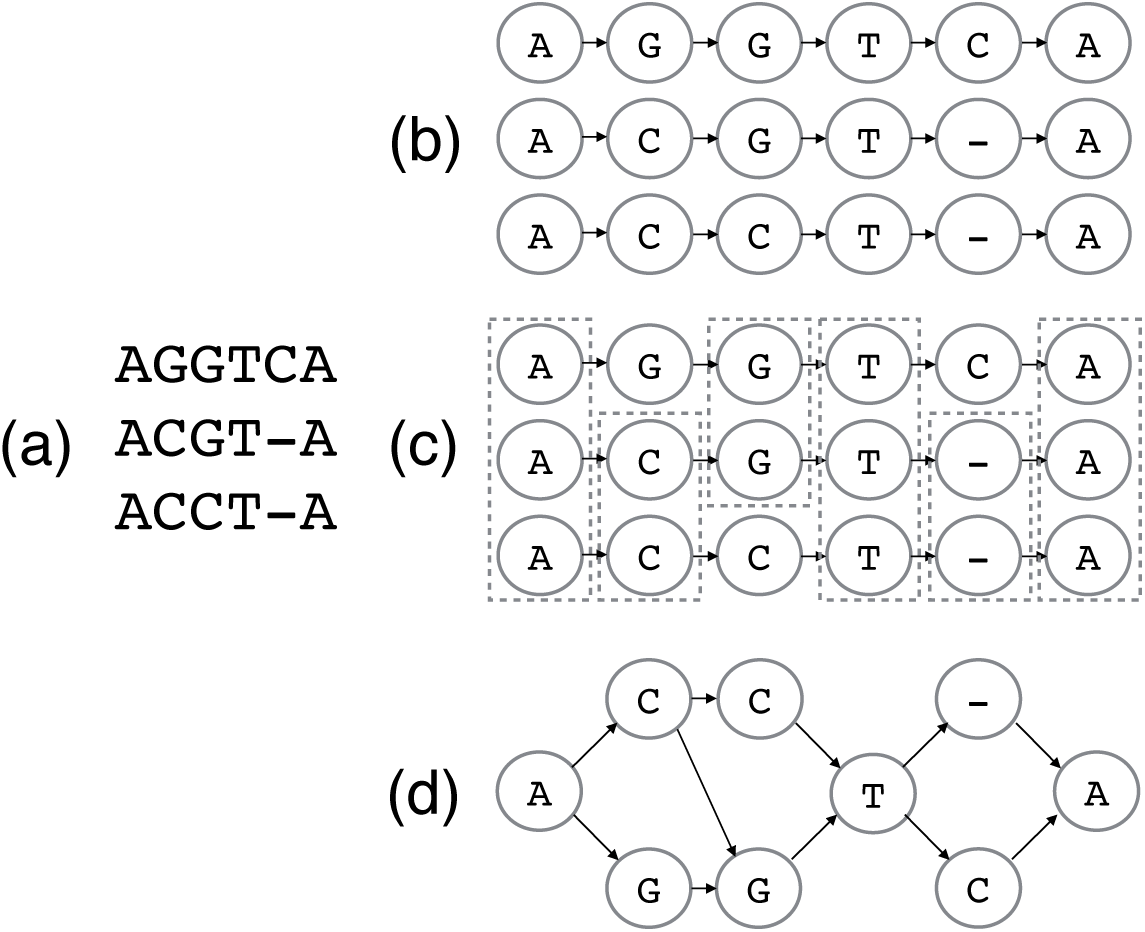
MSA to partial-order graph construction for HLA assembly. Given a pre-computed MSA (a), each sequence is constructed as a chain of verticies connected by directed edges and corresponding positional verticies are aligned vertically (b). For each column, redundant vertices are grouped together (drawn as enclosed in dotted boxes in (c)) and when they are merged, the corresponding partial-order graph (d) is obtained.

### Modification of the HLA-graph via alignment projection

Consider an example novel allele sequence of AGCTCA. It is easy to see that there is no path encoding such allele in the HLA-graph shown in Figure 3(d). In this example, simply adding an edge from the vertex G at column 2 to the vertex C at column 3 is the only modification needed for the graph to include the path that encodes the novel allele. If a novel allele exists in data, there must be sequencing read that contain the differences the nov.el allele has compared to known alleles. Assuming the sequence divergence is small enough for pairwise alignment of the read and a known allele to capture the differences, we can obtain the novel variants. For this reason, we further modify the HLA-graph to include additional paths that encode for novel alleles in a test individual. We achieve this by modifying the previously constructed HLA-graph by projecting the alignments of the reads likely coming from HLA region to known HLA genes.

We first align the extracted reads to the set of reference panel sequences obtained from *M_panel_* using `BWA` (v0.7.15-r1140) [16]. The linear alignments obtained are then projected onto gene-specific partial order graphs. That is if a read is mapped to the HLA-A gene, then the alignment is projected onto the HLA-graph of the gene. Given a read *r*, a subsequence *h* of a known allele *H*, and a pairwise alignment of *r* and *h*, by projection of the alignment to the HLA-graph, our goals are to (1) modify the graph to encode the exact sequence of *r* within the range of columns *h* is encoded in the graph, (2) increment the weight of each edge of the path by 1, and (3) preserve preexisting paths at the same time. When *r* and *h* are identical, the graph must already contain a path that exactly encodes *r* because *H* is in the MSA used to contstruct the graph. When there are few differences such as mismatch, deletion, or insertion identified by the pairwise alignment of *r* and *h*, there are two cases: (1) *r* is already encoded in the graph and (2) *r* is not, thereby requiring modification of graph to encode *r*. For example, ACGTCA does not align perfectly to any of the sequences in Figure 4(a) but it is encoded in the graph as a path. On the other hand, there is no path encoding ACCTGA.

Examples of graph modification by alignment projection are shown in Figure 4. The panel (a) shows a MSA with 3 known alleles and the corresponding POG. Modification for mismatches and deletions are simple because they only require adding a vertex for the mismatched base or a gap (‘-’) symbol. Figure 4(b) illustrates an example where *r* has a deletion of ‘T’ at position 4. A gap vertex is added to the corresponding column and edges are added to connect the newly added vertex to the previous and next base in *r* to obtain a path encoding *r*. Normally, insertion requires a shifting of columns in the MSA/graph because extra columns are required for the inserted bases to be encoded. However, some alignments with insertions do not require a column shifting. An example of an alignment with insertion not requiring a column shifting is shown in Figure 4(c). The read is aligned to *H*_3_ with insertion at position 5 instead of aligning to an allele *H*_1_ with a mismatch at the same position because the alignment score with 1 insertion is higher than the score with 3 mismatches (position 2, 3, and 5 if aligned to *H*_1_). Because of *H*_1_ is in the MSA, the graph already has the column for handling an insertion at this particular column. In this case, we simply insert a vertex with ‘G’ symbol into the corresponding column and connect edges to complete the path for *r*.

Finally, the case of an insertion requiring a shift of columns is depicted as an example in Figure 4(d). The read is aligned to allele *H*_1_ with an insertion of ‘A’ at position 4. To insert a new column between 3rd and 4th columns (also denote them as left and right columns), we first insert a new vertex with a ‘-’ symbol and need to reroute all edges between the left and right columns through the newly inserted gap symbol and redistribute edge weights in order to preserve the preexisting paths. Adjusted weights are shown on the edges in the example. To describe formally, let *L* and *R* be sets of vertices of the left and right columns respectively and *E* be the set of directed edges from *v*_*l*_ ∈ *L* to *v*_*r*_ ∈ *R* with the weight of each edge as *w*({*v*_*l*_, *v*_*r*_}). Additionally, let 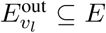 be the set of all outgoing edges of *v*_*l*_ and 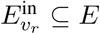 be the set of all incoming edges of *v*_*r*_. Note that there are always 1 or more outgoing edges from *v*_*l*_ and 1 or more incoming edges to *v*_*r*_. After disconnecting all {*v*_*l*_, *v*_*r*_} ∈ *E*, we make a new vertex *v*_*gap*_ with ‘-’ symbol and add an edge {*v*_*l*_, *v*_*gap*_} for each *v*_*l*_ and assign a weight of 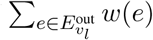. Similarly, we add an edge {*v*_*gap*_, *v*_*r*_} for each *v*_*r*_ with a weight of 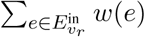. Once the column shifting is done, we can actually process the insertion base exactly same as we handled the case of inserting into a gap column.

**Figure 4.**
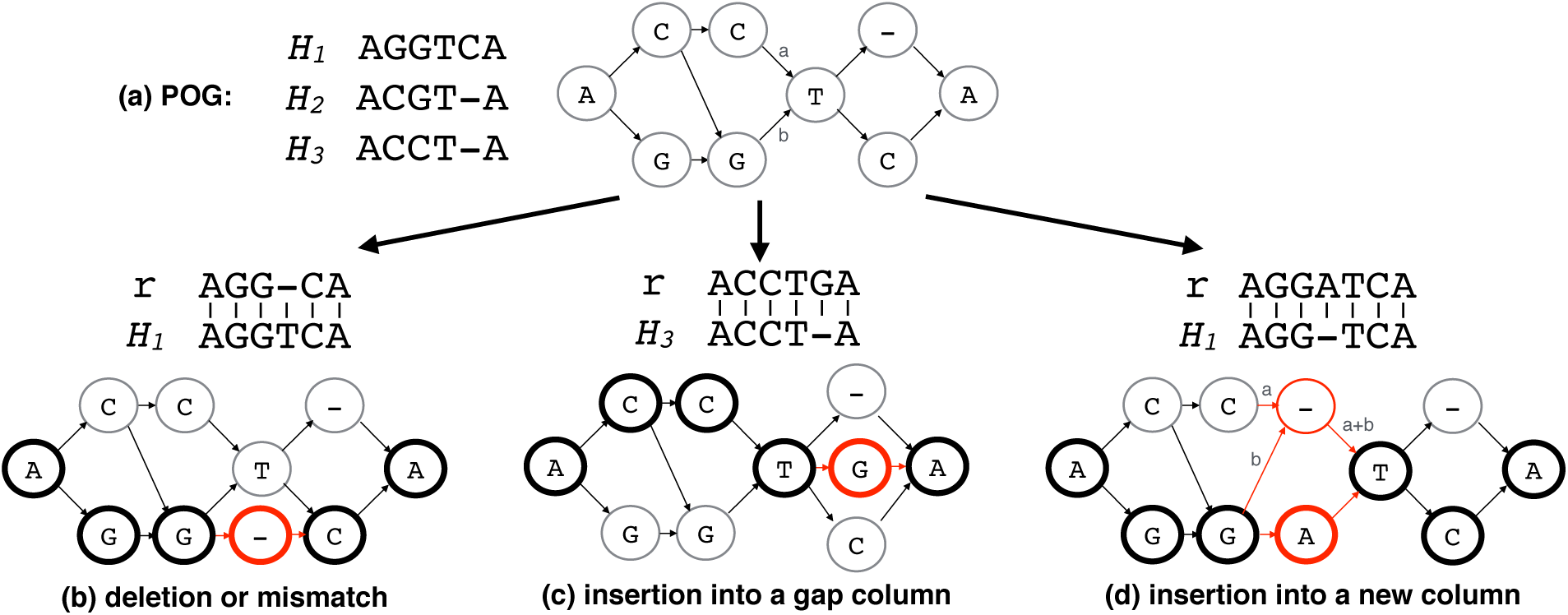
Modification by alignment projection. The same MSA and its corresponding POG from Figure 3 is shown (a). Three examples of the graph modification operations (deletion or mismatch (b), insertion into a gap column (c), and insertion into a new column (d)) are shown respect to the initial POG constructed. For each operation, an alignment of read *r* to one of the known alleles *H_i_* is used to modify the graph. Each operation is applied to the POG and the resulting graph is shown. The nodes and edges that are newly added or changed during the operation is shown in red. The nodes that read path maps are shown as bold circles. For the case of the insertion into a new column, the newly assigned edge weights are explicitly drawn in using *x* and *y* variables.

### Finding paths through the HLA-graph

Given the HLA-graph with weights, assembling HLA alleles can be formulated as the problem of finding two (diploid) paths (they can be identical) that explain the read mapping data (weights and phasing) best. For example, the read depth value for an edge can be thought of as a capacity of the edge in classical flow problems. When considering only the weights, we can find two paths where the sum of the flow values of the paths are maximized. However, this formulation does not handle phasing information embedded by reads or read pairs, therefore it can possibly select erroneous paths that are not consistent with phasing information. For this reason, we want our objective to take both weights as well as phasing information into account. Since read information is embedded on the HLA-graph, we can check if two neighboring variant sites can be phased directly by a read or read pair. For example, given two heterozygous sites with A/T and G/C, a read or a read pair carrying ‘A’ followed by ‘G’ at these sites indicates the chromosomal phase of ‘AG’ since the sequencing read is assumed to come from a contiguous segment in a chromosome. In our method, we first investigate variant regions individually to select locally phased paths with strong read support and construct a set of full-length paths through the HLA-graph by connecting the locally phased paths that can be further phased by read or read pair. Each of these full-length paths is considered as a candidate allele and the best pair among the candidates with maximum read and phasing support is selected as the output. To only consider nonzero-weight full-length paths, we remove all zero-weight edges and disconnected vertices prior to finding paths.

### HLA-graph to bubble graph

We first focus on the parts of the HLA-graph where variations are captured, which are often referred to as bubbles in sequence assembly graphs [20, 21, 22, 23]. In the HLA-graph, we define a *bubble* as a region (3 or more consecutive columns) where there is only one vertex each in the leftmost and rightmost columns and the rest of the columns must have 2 or more vertices. Let *s* and *t* be the vertices in the leftmost and the rightmost columns respectively. Any vertex in the bubble is reachable from *s* and one or more paths exists from any vertex in the bubble to *t*. Any two distinct paths that goes through a bubble must go through *s* and *t*. Bubbles naturally capture varying sites between two alleles in the graph. The regions that are enclosed by dotted line in Figure 5(a) are examples of bubbles. On the other hand, a region that is completely shared by all paths through the HLA-graph represents a conserved region. Without the loss of generality, the HLA-graph can then be thought of as a chain of bubbles, where two neighboring bubbles are connected by a linear path of length 0 or longer (Figure 5). For simplicity, we can connect the bubbles without the linear paths as they do not play any role in determining the phase of a haplotype. We call this a bubble graph and bubbles can easily be recognized in the HLA-graph because of its structure.

**Figure 5.**
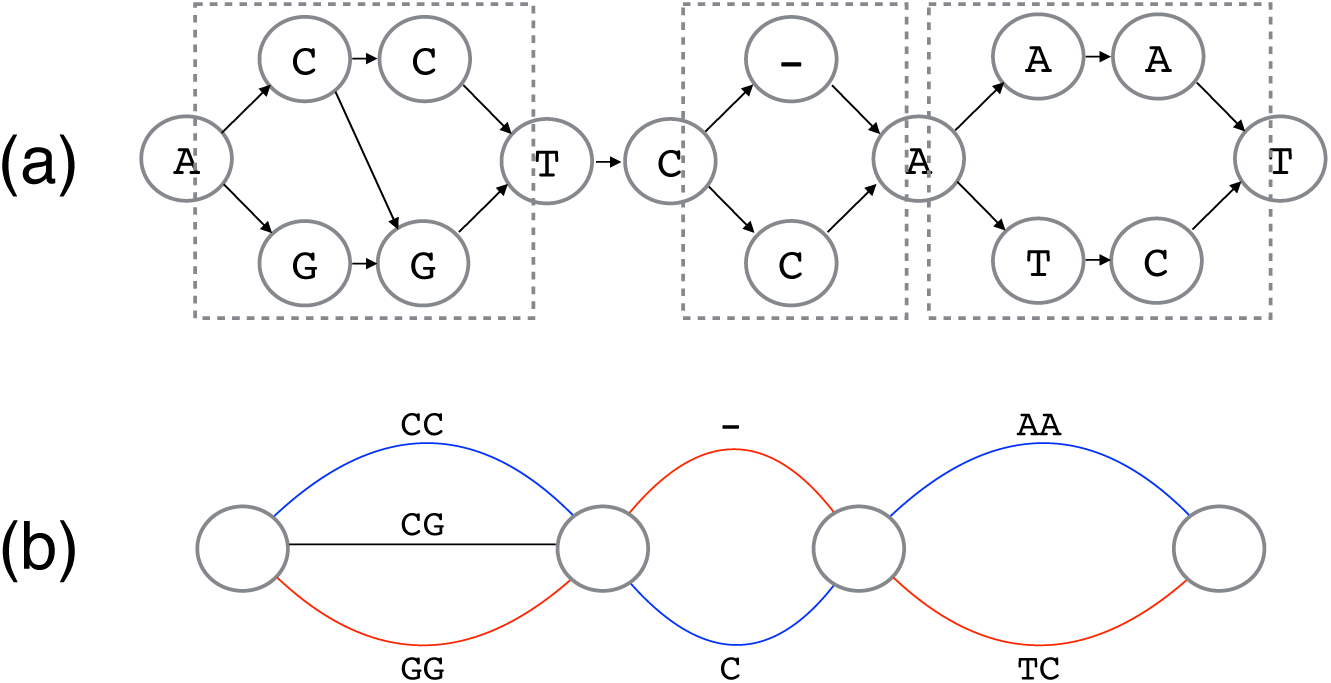
HLA-graph to bubble graph. An example of HLA graph with 3 bubbles (enclosed in dotted boxes) are shown (a) and its corresponding bubble graph is shown (b). Best paths through the bubbles can be thought of as a pair of distinct colored paths (shown in red and blue).

### Finding the best set of paths in a bubble

Ideally, we want to find exactly 2 paths per bubble since the ploidy number is 2 for humans. However, bubbles may contain more than 2 paths because of sequencing errors or misalignment. Therefore, we first identify all paths that are phased by a read or read pair. For each bubble, we can use a modified breadth first search (BFS) technique to obtain all paths that go through the bubble. In order to avoid enumerating over all paths through a bubble, we prune any path without a read backing the sequence encoded by the path at each iteration of BFS. For a path in the bubble to be retained, it must be supported by at least one read phasing the entire path. We can simply compute the set of phased reads for a path by taking a series of intersections of read sets maintained by each edge in the path. Each phased path through a bubble is a called a *bubble path*. Given multiple bubble paths from a bubble, our goal is to select the best pair of paths. We iterate over all possible pairs of bubble paths to calculate the posterior probability of each pair given all reads aligned to the bubble to find the pair that gives the maximum probability. We write the posterior probability of a given genotype as

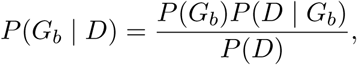

where *G*_*b*_ is a genotype and *D* is the alignments of all reads aligned over the bubble. The genotype is a pair of bubble paths *G*_*b*_ = (*H*_*b*1_, *H*_*b*2_). Each *d* ∈ *D* is an alignment string of a segment of a read and *d*^*i*^ is the *i*-th symbol in segment *d*. Similarly, 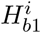 is the *i*-th symbol in *H*_*b*1_. *P(D)* is constant and we assume that the prior probability *P*(*G*_*b*_ = (*H*_*b*1_, *H*_*b*2_)), is uniformly distributed over all genotypes. We can then compute the conditional probability *P*(*D*│*G*_*b*_) by adopting widely used formulations [18, 24] with small variations to allow multiple positions and base ‘N’ case that can be present from sequence data. We iterate over each read and compute *P*(*D*│*G*_*b*_) as a product of the conditional probability of each read *d*. Since a read must come from one of the two chromosomes, and we assume that *d* is equally likely to come from either one of them, we can rewrite it as a sum of average of two cases where *d* is from *H*_*b*1_ and *H*_*b*2_ and it is 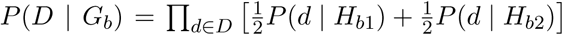.To compute the conditional probability of each *d* given a bubble path *H*_*b*_, we iterate over each pair of corresponding positions *d*^*i*^ and 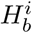jointly, assuming each *d*^*i*^ is conditionally independent of each other given 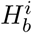. Therefore, the probability of each base *d*^*i*^ given a pair of corresponding genotype bases 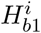 and 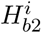 is 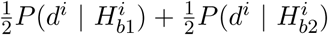. The probability of seeing a base given an allele is defined as

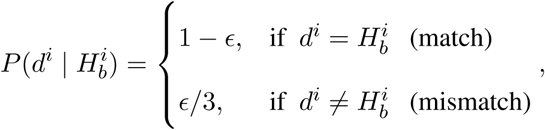

where є of base symbol *d*^*i*^ is the error probability obtained from the phred score of the base. For the case of *d*^*i*^=‘N’, we simply estimate the probability as 1/4. Instead of selecting *H*_*b*_ from all possible │*d*│-mers, we limit to only the bubble paths found in the bubble and iterate over all pairs to select a pair of bubble paths *P*_*b*_ that jointly gives the maximum probability:

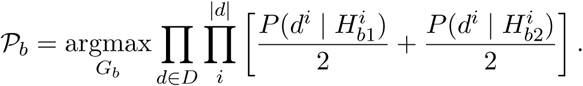

### Phasing paths

We now have an ordered list of bubbles, and a list of “best” read-backed phased bubble paths for each bubble. The goal here is to find a set of candidate paths through all the bubbles by merging one bubble at a time iteratively from left to right, connecting bubble paths that are phased by a read or read pair. Two paths are said to be phase-consistent if there is a read or read pair spanning both paths. This can be checked easily by taking an intersection since each bubble path maintains a set of phasing reads. Given a set of already merged bubble paths *Ƥ*_*m*_ from the first *i* − 1 bubbles and a set of bubble paths *Ƥ*_*b*_*i*__ from the *i*-th bubble, we look at all pairs of paths *Ƥ*_*m*_ × *Ƥ*_*b*_*i*__ and keep only pairs that are phase-consistent and connecting each of such pairs as one path. We also update the phasing-read set for each merged path.

### Selecting the best pair of candidate alleles

Once the assembly by bubble merging is finished, we have a set of merged bubble paths through all bubbles. By placing back the linear chains that were ignored during bubble merging to original positions (in between bubbles), we have a full-length candidate allele *H*_*i*_ for each merged bubble path. Let *C* be the set of all candidate alleles and *B* be a set of all bubbles. Our goal is to select a pair of alleles (*H*_1_, *H*_2_) ∈ *C* × *C* that has the most consistent phasing support over all bubbles. We first define a scoring metric that checks for strength of phasing support jointly for a pair of allele *H*_1_ and *H*_2_ between a pair of consecutive bubbles *b*_*i*_ and *b*_*i*+1_ and it is defined as

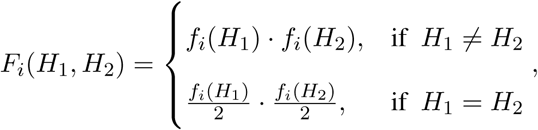

where *f*_*i*_(*H*)is the inter-bubble phasing fraction. The fraction *f*_*i*_(*H*) is the ratio of the number of phasing reads for allele *H* between *b*_*i*_ and *b*_*i*+1_ and the number of total phasing reads. When considering two paths at the same time, there can be regions where the paths overlap (homozygous: shown as purple edges in Figure 2(e)) and separate (heterozygous: shown as blue or red edges in Figure 2(e)). For homozygous sections of the paths, that is *H*_1_ = *H*_2_, *f*_i_(*H*) is halved to keep balance between calling homozygous and heterozygous alleles. We can calculate a product of *F*_*i*_ over all pairs of neighboring bubbles to check the consistency of phasing support for the pair. Finally, we select the pair of alleles Ƥ that maximizes the product over all pairs of alleles:

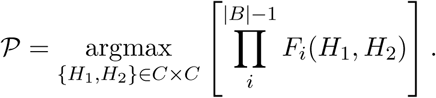

## Description of data used for evaluation

### Simulated data

For each of the 6 HLA genes (HLA-A, HLA-B, HLA-C, HLA-DQA1, HLA-DQB1, and HLA-DRB1) tested, we randomly selected 2 full-length alleles from the IPD-IMGT/HLA database (v3.24.0) and repeated this for 100 replicates, resulting in a total of 200 alleles to simulate. The exact number of full-length alleles in the database is reported in Table 1. For each replicate, we simulated 25X coverage of paired-end WGS data for each allele, making it 50X coverage for each locus. For the simulation of paired-end reads, we used an Illumina read simulator, `pIRS` [25], which simulates using empirical base-calling and GC%-depth profiles trained from Illumina re-sequencing of known samples. We used 2 × 100bp for the read length and 500 +/− 50bp for the mean and the standard deviation of the insert size.

### Illumina Platinum Genomes

Illumina has sequenced 17 individuals (CEPH/Utah pedigree 1463) in a three generation family using their high-coverage PCR-free paired-end WGS assay (2 × 101bp). These genomes are often referred to as the Illumina Platinum Genomes [26]. The family pedigree is shown in Figure 6. Many individuals in this family are extensively investigated by the genomics community especially NA12891-NA12892-NA12878 trio as well as NA12898-NA12890-NA12877 trio. The read alignments to the GRCh37 version of the Human genome for all 17 individuals were downloaded from the Illumina Platinum Genomes page hosted on Google Cloud Platform (Table S2 in Supplementary Materials) and they were realigned to GRCh38 version of the Human genome.

**Figure 6.**
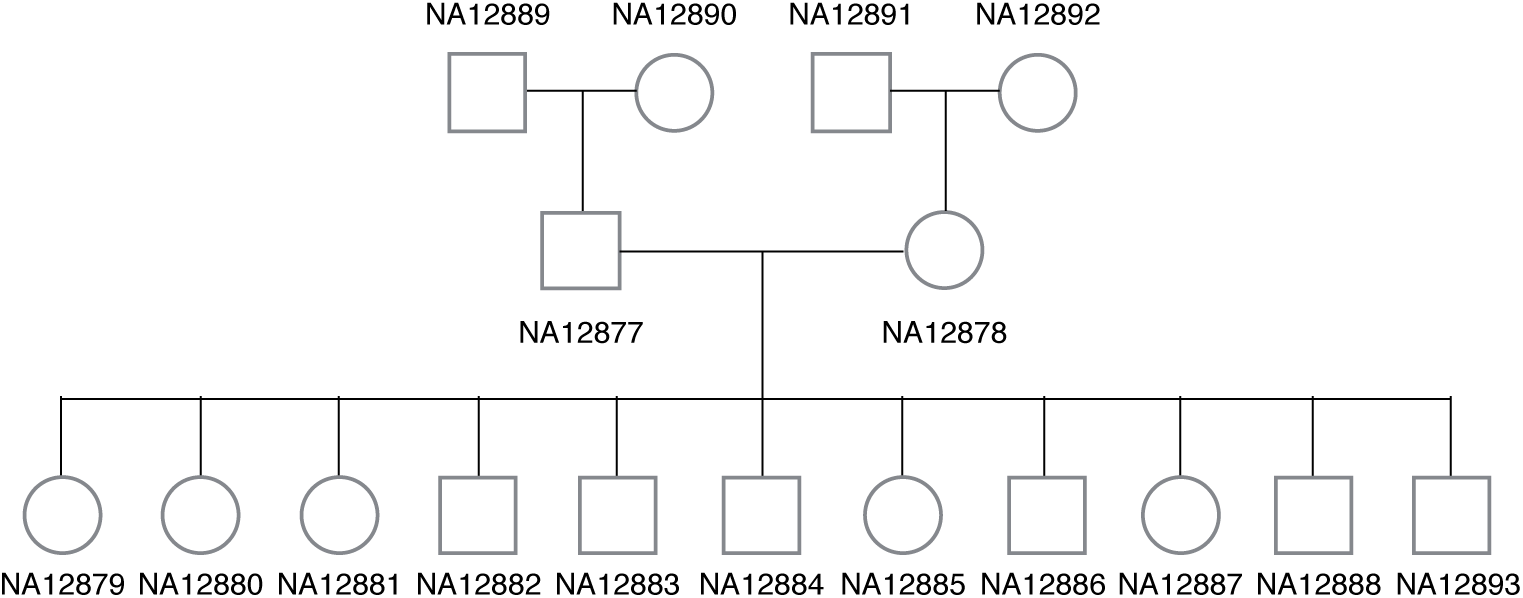
CEPH/Utah pedigree 1463. The family pedigree of Illumina platinum genomes is shown.

### 1000 Genomes

The 1000 Genomes Project [27] has produced various personal genomic data. Among these, there are 11 individuals whose high-coverage WGS sequencing data along with validated HLA typing results [28] are available. This dataset covers a wide ethnic diversity (1 Colombian from Medellín, 3 Utah residents with Northern and Western European ancestry in a trio, 1 Japanese from Tokyo, Japan, 3 Yoruban from Ibadan, Nigeria in a trio, 1 person of African ancestry from the southwestern United States, 1 person of Mexican ancestry from Los Angeles, and 1 Toscani from Italy) covering various different HLA types, making it an ideal dataset to test on. The bam files aligned to the GRCh38 version of the Human genome were downloaded from the 1000 Genomes data portal (http://www.internationalgenome.org/data-portal). For the Utah resident trio and the Yoruban trio, we downloaded fastq files and realigned to the GRCh38 version because GRCh38 bam files were not available. The complete list of links to the data downloaded is in Table S2 in Supplementary Materials.

## Results

### Simulation

In order to check that our method performs well, we tested our method on simulated data (see “Materials and Methods”). For each of the 6 HLA genes, 2 alleles from the set of full-length gene sequence in the IPD-IMGT/HLA database were randomly chosen. We repeated for a total of 100 replicates, resulting in 200 randomly selected alleles across all replicates. For each replicate, we simulated 50X coverage (25X for each haplotype) of paired-end WGS data. We compared `Kourami`, `PHLAT`, and `HLA*PRG` on the simulated data. Our method was evaluated using all 1200 alleles (2 alleles × 6 genes × 100 replicates), however, not all alleles could be used for the evaluation of `PHLAT` and `HLA*PRG` as both tools use their own digested format of the HLA database and built it into the tools so that the content of the database cannot be updated by a user. The database versions used by `PHLAT` and `HLA*PRG` are older compared to the version (v3.24.0) used for `Kourami`. Given a set of WGS data of an individual with an allele that is not in the database built into `PHLAT` and `HLA*PRG` both tools will fail to type the allele correctly as they are designed strictly to find the nearest match among the known alleles. For this reason, evaluation of `PHLAT` and `HLA*PRG` are only based on the subset of simulated alleles (1011 for `PHLAT` and 990 for `HLA*PRG`) that are in the database versions they use. For `PHLAT`, 4-digit ‘P’ resolution was used and 6-digit ‘G’ resolution was used for `HLA*PRG` and `Kourami` for evaluation. Table 2 shows the number of correctly inferred alleles as well as the accuracy for each HLA gene tested. For our method, we report both the typing and assembly accuracy where an assembled allele is correct if the output sequence is identical to the true allele sequence (no mismatch or indel). Even when an assembled allele is not identical to its expected true sequence, the typing of the allele may be correct if the closest sequence (minimum edit distance) in the database is the true allele. `PHLAT` achieves 93.6% across all HLA genes tested (89.8% for Class I and 96.6% for class II). `HLA*PRG` and our method perform equally well, achieving 99.8% typing accuracy across all genes (99.5% for class I and 100% for class II). Additionally, `Kourami` achieves 99.3% assembly accuracy.

**Table 2:** HLA typing performance on simulated data.

|  | Class I |  |  | Class II |  |  |
| --- | --- | --- | --- | --- | --- | --- |
|  | A | B | C | DQA1 | DQB1 | DRB1 |
| PHLAT (4-digit ‘P’) | 0.85 (147/172) | 0.91 (135/149) | 0.95 (122/129) | 0.93 (178/191) | 0.98 (169/173) | 0.99 (195/197) |
| HLA*PRG (6-digit ‘G’) | 1.00 (174/174) | 0.99 (143/144) | 0.99 (115/116) | 1.00 (197/197) | 1.00 (159/159) | 1.00 (200/200) |
| Kourami (Type) | 1.00 (199/200) | 1.00 (200/200) | 0.99 (198/200) | 1.00 (200/200) | 1.00 (200/200) | 1.00 (200/200) |
| Kourami (Sequence) | 0.99 (198/200) | 1.00 (200/200) | 0.98 (195/200) | 1.00 (200/200) | 1.00 (199/200) | 1.00 (200/200) |

Accuracy is shown as a fractional number and the fraction of number of correctly typed alleles and total number of alleles tested is shown in parenthesis.

### Novel Allele Detection

The major benefit of our method is that it can assemble novel alleles across the typing exons, therefore its typing capacity is not limited by known alleles as is the case with other database-matching methods. Unlike the database-matching methods, `Kourami` uses the known alleles in the input database only to construct the HLA-graph that serves as a template for reference-based assembly but does not discriminate between the paths that encodes known alleles and novel alleles.

In order to demonstrate the ability to assemble novel alleles, we evaluated `Kourami` across various data where ground truth is known. We tested on the simulated data and the real data with previously validated HLA types (NA12878-NA12891-NA12892 Platinum trio and 11 samples from the 1000 Genomes Project) with a modified database of known alleles so that `Kourami` is not aware of true allele sequences. For each sample, we randomly selected one allele from each of the 6 HLA genes and removed the selected alleles from the reference MSAs (full-length and exon-only) provided by the release 3.24.0 of the IPD/IMGT-HLA database. When removing an allele, we removed all entries in the ‘G’ group that the allele belongs to and the entire list of alleles removed from each individual is shown in Tables S3, S4, and S5 in Supplementary Materials. We removed corresponding rows for the alleles from *M_gene_* and *M_coding_* and generated new 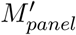. The number of ‘G’ group alleles removed is 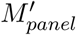 and the bam files obtained were used as inputs to `Kourami`. Note that this experiment cannot be done with `PHLAT` and `HLA*PRG` as the database of known alleles are built into the tools.

`Kourami` correctly assembled 98.3%, 100%, and 98.3% of the removed alleles for simulation data, the Platinum trio, and 11 samples from the 1000 Genomes Project respectively (Table 3). Among 1000 Genomes samples, the only incorrectly assembled allele (supposed to be B*38:01:01) had 1 base-pair difference to the correct sequence. When the 59 correctly assembled allele sequences are aligned to all alleles in 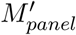, many alleles were aligned equally well to a large number of known alleles. For example, C*05:01:01 alleles aligned to 122 other alleles with just 1 base-pair substitution. Among them, a significant portion contained base differences that result in protein-coding changes in typing exons. This shows that the databasematching methods such as `PHLAT` and `HLA*PRG` are not only unable to discover novel alleles but also faced with a problem of selecting the best out of many alleles with equally similar sequences.

**Table 3:** Novel allele recovery.

|  | Simulation | Platinum trio | 1000 Genomes |
| --- | --- | --- | --- |
| # removed alleles | 596 | 15 | 60 |
| # recovered alleles | 586 | 15 | 59 |
| % recovered alleles | 98.3 | 100 | 98.3 |

## Illumina Platinum Genomes

### Platinum trio with validated results

Among the Illumina Platinum Genomes, we first ran `Kourami`, `PHLAT`, and `HLA*PRG` on the trio (NA12891, NA12878, and NA12892) with the previously validated 4-digit HLA types for 6 HLA genes (HLA-A, HLAB, HLA-C, HLA-DQA1, HLA-DQB1, HLA-DRB1) [29]. `Kourami` and `HLA*PRG` perfectly called the correct types where `PHLAT` missed a call in the HLA-C gene in NA12891. In a previously published article [12], `PHLAT` called all 12 alleles correctly and the difference may be due to the fact that in our evaluation all software were run on the set of reads that aligned to xMHC/HLA region of chromosome 6 and unmapped reads. Extraction of subset of reads by read mapping location and including unmapped reads are common practice to reduce computational time, and a similar technique was used in [9].

### Trio consistency and inferred haplotypes

The pedigree of Illumina platinum genomes include many third generation offspring and only the top right-hand trio in Figure 6 has previously validated HLA typing results. Since this trio includes the mother (NA12878) of all third generation offspring, if HLA typing results are trio-consistent across all trios and all second-generation-haplotypes are present in one of the children, we can theoretically infer HLA haplotypes of the second-generation male (NA12877) as well as the half of HLA haplotypes in the first-generation individuals (NA12889 and NA12890).

We tested all 3 methods to determine whether predictions are trio-consistent across all trios (trio consistency shown in Table 4). `Kourami` and `HLA*PRG` agreed on all 204 alleles at 6-digit ‘G’ resolution and the predicted alleles were trio-consistent and inferred haplotypes across HLA genes (intra-gene phased) are shown in Figure 7. `PHLAT`’s predictions were trio-consistent only for HLA-C and HLA-DQB1 when evaluated at 4-digit ‘P’ resolution, and additionally for HLA-A when evaluated at 2-digit resolution. Although, we do not know the true HLA types for the rest of 14 individuals, it is very likely that the predicted HLA types are correct given that all typing results are consistent. Low trio-consistency ratios for `PHLAT` in Table 4 is mainly due to mistyped alleles in HLA-A and HLA-B in the NA12877 individual. Assuming the predicted HLA types for the pedigree are correct, no recombination seems to have occurred, leaving no disruption in ancestral haplotypes. In Figure 7, we labeled the haplotypes that are originating from the first generation members as paternal-grand-father1/2 (PGF1, PGF2), paternal-grand-mother1/2 (PGM1, PGM2), maternal-grand-father1/2 (MGF1, MGF2), and maternal-grand-mother1/2 (MGM1, MGM2). The haplotypes that are passed to second generation individuals are numbered ‘1’ to keep the numbering consistent in the 3rd-generation. Among 11 third-generation offspring, all 4 possible pairs of haplotypes were observed (2 PGF+MGF, 2 PGF+MGM, 4 PGM+MGF and 3 PGM+MGM).

**Table 4:** Trio consistency over 12 trios in platinum genomes.

|  | Class I |  |  | Class II |  |  |
| --- | --- | --- | --- | --- | --- | --- |
|  | A | B | C | DQA1 | DQB1 | DRB1 |
| PHLAT (2-digit) | 1.0 (24/24) | 0.71 (17/24) | 1.0 (24/24) | 0.79 (19/24) | 1.00 (24/24) | 0.96 (23/24) |
| PHLAT (4-digit 'P') | 0.67 (16/24) | 0.42 (10/24) | 1.0 (24/24) | 0.79 (19/24) | 1.0 (24/24) | 0.46 (11/24) |
| HLA*PRG (6-digit 'G') | 1.00 (24/24) | 1.00 (24/24) | 1.00 (24/24) | 1.00 (24/24) | 1.00 (24/24) | 1.00 (24/24) |
| Kourami | 1.00 (24/24) | 1.00 (24/24) | 1.00 (24/24) | 1.00 (24/24) | 1.00 (24/24) | 1.00 (24/24) |

Trio consistency is shown as a fractional number and number of alleles consistent is shown as a fraction in parenthesis

**Figure 7.**
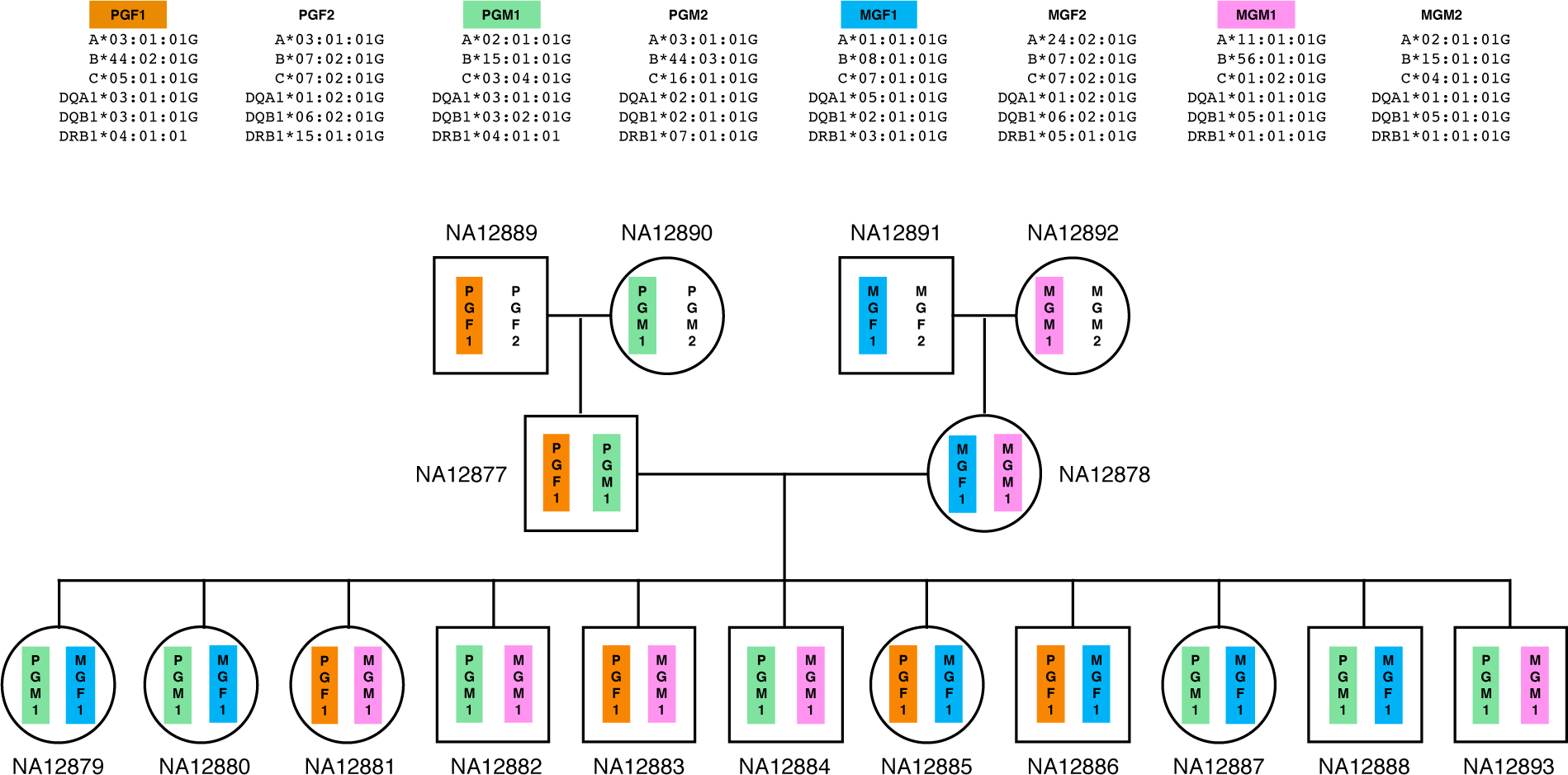
HLA haplotypes in Illumina Platinum pedigree. This picture shows Illumina platinum pedigree with the predicted HLA haplotype information. Four haplotypes (PGF1, PGM1, MGF1, and MGM1) found in second generation are intermixed in third generation offsrping. Note that only the haplotypes that are passed down to 2nd and 3rd generations are colored. A haplotype drawn on the left is always inherited from his/her father. For the first generation, this information is missing and the haplotypes that are passed to next generation are arbitrarily drawn on the left.

## 1000 Genomes

We tested all three methods on this data set and the result is summarized in Table 5. `PHLAT` called 93 out of 122 alleles correctly, resulting in 76% accuracy when evaluated at 4-digit ‘P’ resolution, and 89% when evaluated at 2-digit resolution. `HLA*PRG` results were consistent with what was previously reported [12], resulting in 1 error (99.2% accuracy). Our method correctly called all of the alleles without any differences in bases. Note that the total number of alleles tested for DQA1 is 12 instead of 22 (2 alleles x 11 individuals) because the validation data for 1000 genomes [28] does not report DQA1 types. DQA1 type validation is only available for 6 individuals [29].

**Table 5:** HLA typing performance on 11 individuals from 1000 Genomes project.

|  | Class I |  |  | Class II |  |  |
| --- | --- | --- | --- | --- | --- | --- |
|  | A | B | C | DQA1 | DQB1 | DRB1 |
| PHLAT (2-digit) | 0.82 (18/22) | 0.82 (18/22) | 0.91 (20/22) | 0.83 (10/12) | 1.00 (22/22) | 0.95 (21/22) |
| PHLAT (4-digit 'P') | 0.68 (15/22) | 0.55 (12/22) | 0.77 (17/22) | 0.83 (10/12) | 0.95 (21/22) | 0.82 (18/22) |
| HLA*PRG (6-digit 'G') | 1.00 (22/22) | 1.00 (22/22) | 1.00 (22/22) | 1.00 (12/12) | 1.00 (22/22) | 0.95 (21/22) |
| Kourami (6-digit 'G') | 1.00 (22/22) | 1.00 (22/22) | 1.00 (22/22) | 1.00 (12/12) | 1.00 (22/22) | 1.00 (22/22) |

Accuracy is shown as a fractional number and the fraction of number of correctly typed alleles and total number of alleles tested is shown in parenthesis.

## CPU and memory usage

`Kourami` is able to assemble and type HLA alleles given WGS data in a fraction of the time compared to the state-of-art methods such as `PHLAT` and `HLA*PRG` with a moderate use of memory. We compared the CPU and memory usage using the WGS of NA12878 from Platinum Genomes data (2 x 101bp 55x). All tests were run on the input of the reads aligning to xMHC region and unmapped reads. `HLA*PRG` was the slowest–taking 54.62 CPU hours, while `PHLAT` took 10.73 CPU hours and `Kourami` only took 0.09 CPU hours (611x speedup compared to `HLA*PRG`). `HLA*PRG` required the most amount of memory–consuming peak memory of 78.9 Gbytes, while `PHLAT` and `Kourami` used 3.6 Gbytes and 4.3 Gbytes respectively. `HLA*PRG` requires many more CPU hours and a larger amount of memory usuage because of the expensive dynamic-programming-based alignment to graph. `Kourami` relies on fast NGS aligners to align reads against known alleles first and project obtained alignment to HLA-graph to significantly reduce the computational time without sacrificing assembly or typing accuracy.

## Discussion

We have shown that our HLA assembly method can accurately reconstruct both haplotypes that span the typing exons of HLA genes by using a graph representation of known alleles as a guide, and the produced haplotype sequences can be used to successfully carry out HLA typing given high coverage (> 30-fold) paired-end WGS. WGS carried out for other analysis can be used to type individual’s HLA types without the use of another experiment (SBT and other molecular assays).

Most notably, the ability to discover rare and novel alleles is achieved by taking an advantage of the flexibility of POG, combined with graph modification and it is instrumental in both research and clinical settings. It is important to note that previously available computational methods using non-targeted sequencing data cannot discover novel alleles because they are designed to find the best matching allele among the known alleles. Especially with continuously decreasing cost of obtaining a personal genome, personal WGS will only become more widely available, and our method can deliver accurate HLA typing without additional experiments and cost. Also, `Kourami` is able to assemble and type at 6 digit ‘G’ resolution at a fraction of the time compared to other methods with a moderate amount of memory usuage.

One limitation of our method is that it only supports WGS as it needs enough reads to cover both haplotypes for each typing locus, and does not work on other NGS assays, such as WES or RNA-Seq. Since WES is being used widely, as the cost for WES is lower compared to that of WGS, it is useful to be able to type HLA genes using WES. However our testing (not shown) shows that it is difficult to accurately assemble a full length sequence across the typing exons with WES because there are regions for which no reads are sequenced. This may be due to biases that WES has been reported to have [30] as well as decrease in effectiveness in detecting variants when using WES compared to WGS [30, 31]. Additionaly, high-coverage WGS is required to ensure accurate HLA assembly or typing. We randomly sampled coverages of 20x, 30x, and 40x from NA12878 data (Illumina Platinum Genomes) for 5 replicates and tested `Kourami` on these samples. Assembly and typing stays accurate down to 30x coverage (accuracy of 0.97 across the HLA genes) but at 20x coverage, the accuracy drops to 0.82 (Supplementary Table S6). This should not be a surprise as haplotype-resolved assemblies of human genomes used ≈100x coverage of NGS data [32, 33].

Highly accurate results from our method signifies the recent advancement in handling genetic variation using graph structures to encode variations found in multiple reference genomes [34, 35, 36, 13]. Specifically in `Kourami`, the minimal representation of POG allows the consistent graph modification via alignment projection and this in turn enables capturing of novel alleles as paths through the graph. At the same time, it reduces computational time greatly without scarificing accuracy, and this can be beneficial when used in high-demand clinical settings. Our approach can also be extended as a general method of using graph structures as guide to reference-based assembly of high diversity gene families.

## Availability and Implementation

`Kourami` is open source and freely available at https://github.com/Kingsford-Group/Kourami. It is implemented in Java and supported on Linux, Mac OS X, and Windows.

## Acknowledgements

We thank S. Kim of Illumina for helping us in the early stage of this research. We would also like to thank D. DeBlasio, C. Ma, G. Marçais, N. Sauerwald, M. Shao, B. Solomon, T. Wall, H. Wang, and H. Xin for valuable discussions and comments on the manuscript. This research was funded in part by the Gordon and Betty Moore Foundation’s Data-Driven Discovery Initiative through Grant GBMF4554 to C.K., by the US National Science Foundation (CCF-1256087, CCF-1319998) and by the US National Institute of Health (R01HG007104).

---

**S1.**
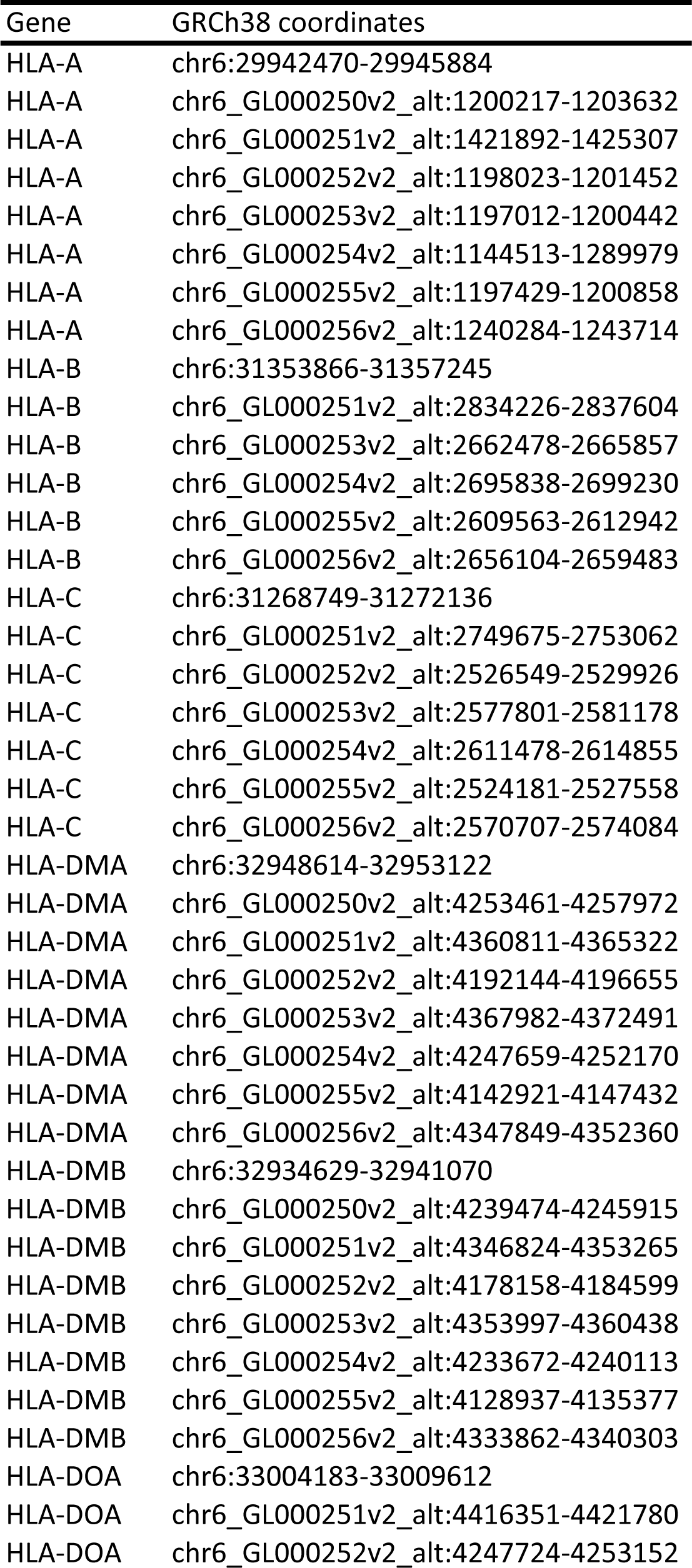
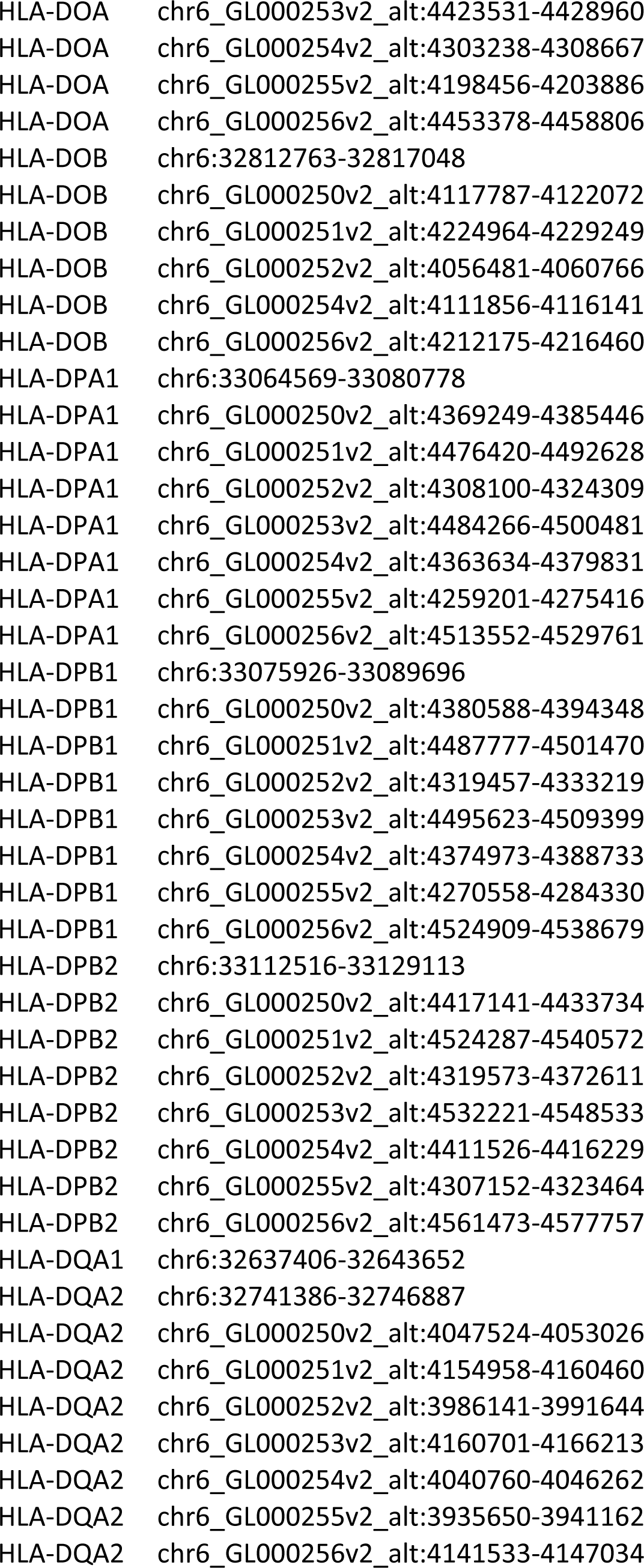
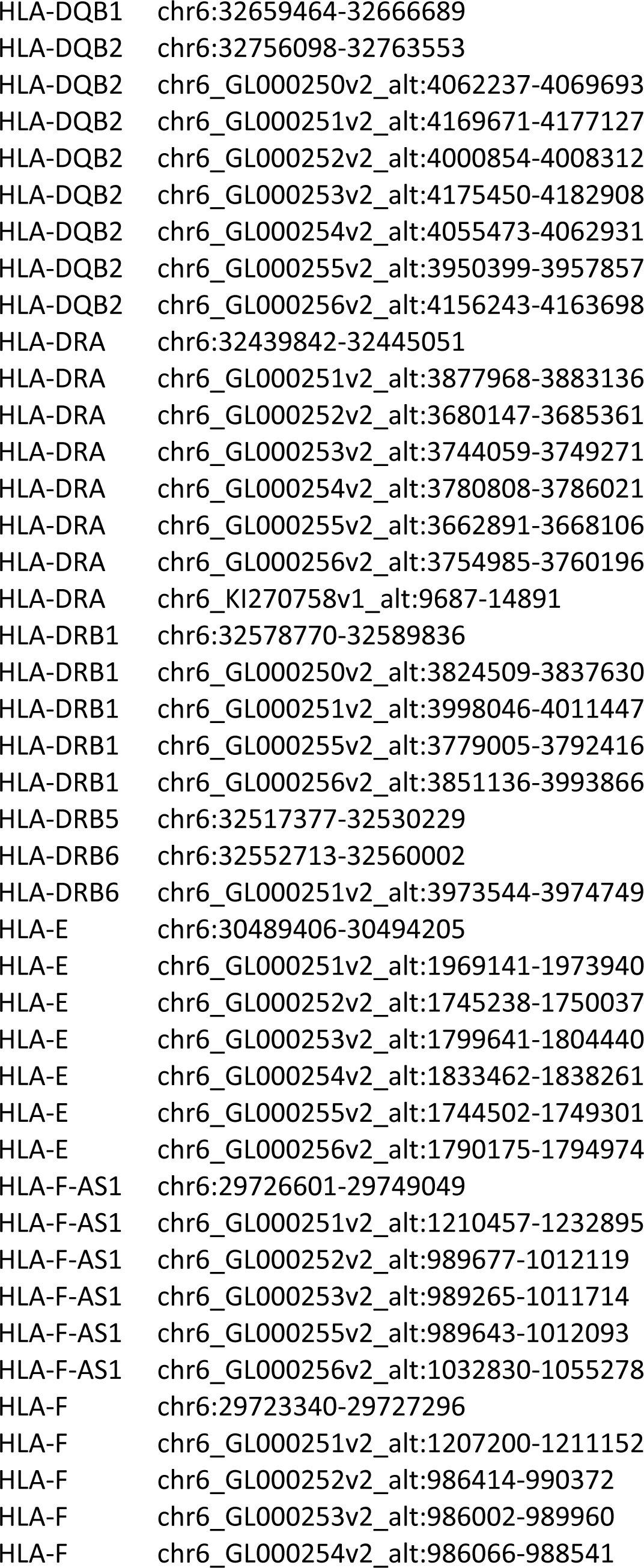
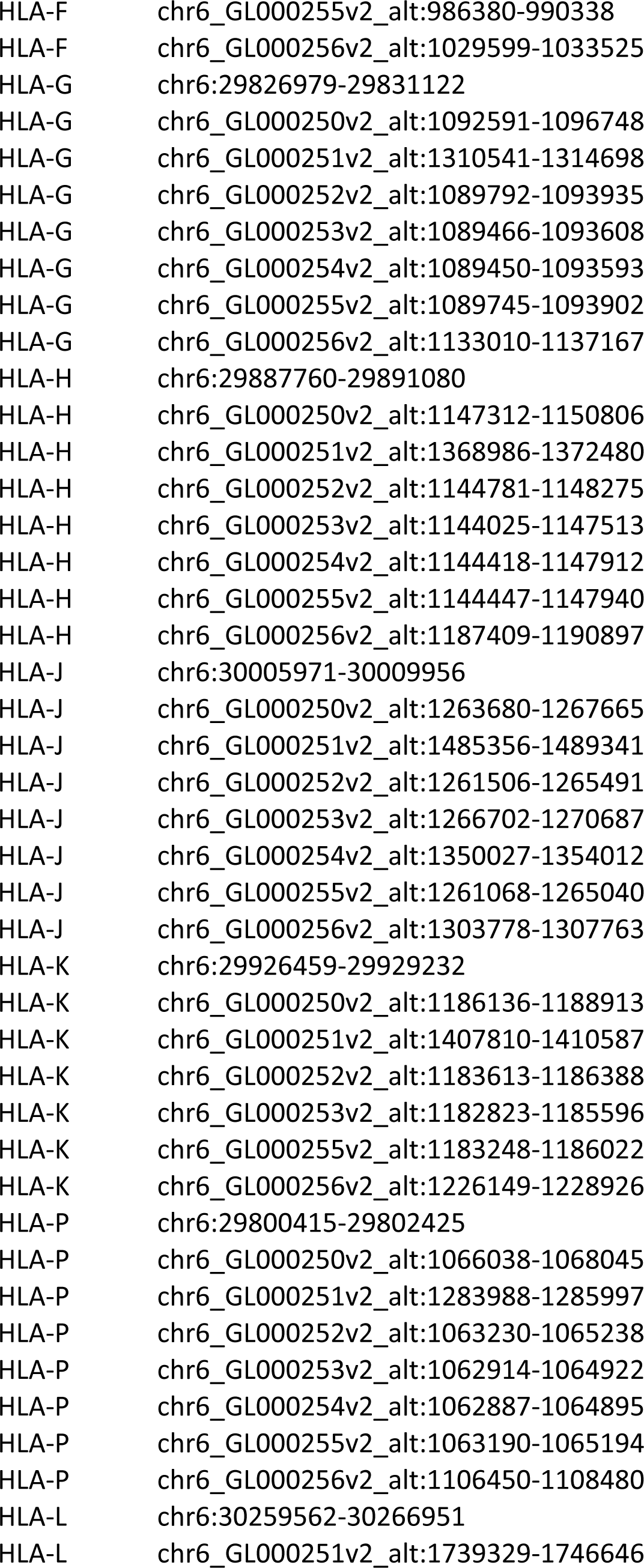
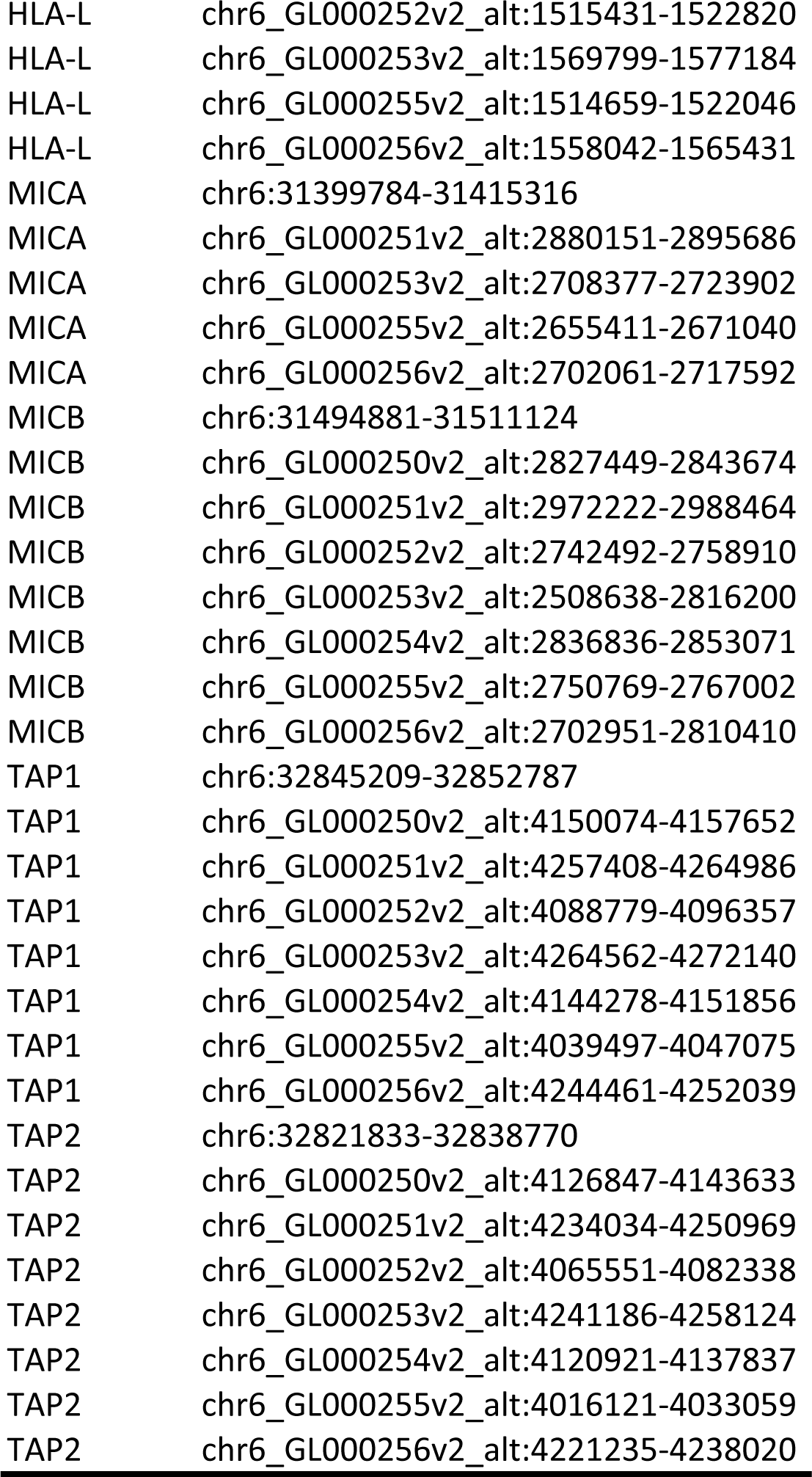
GRCh38 coordinates of HLA genes.

**S2.**
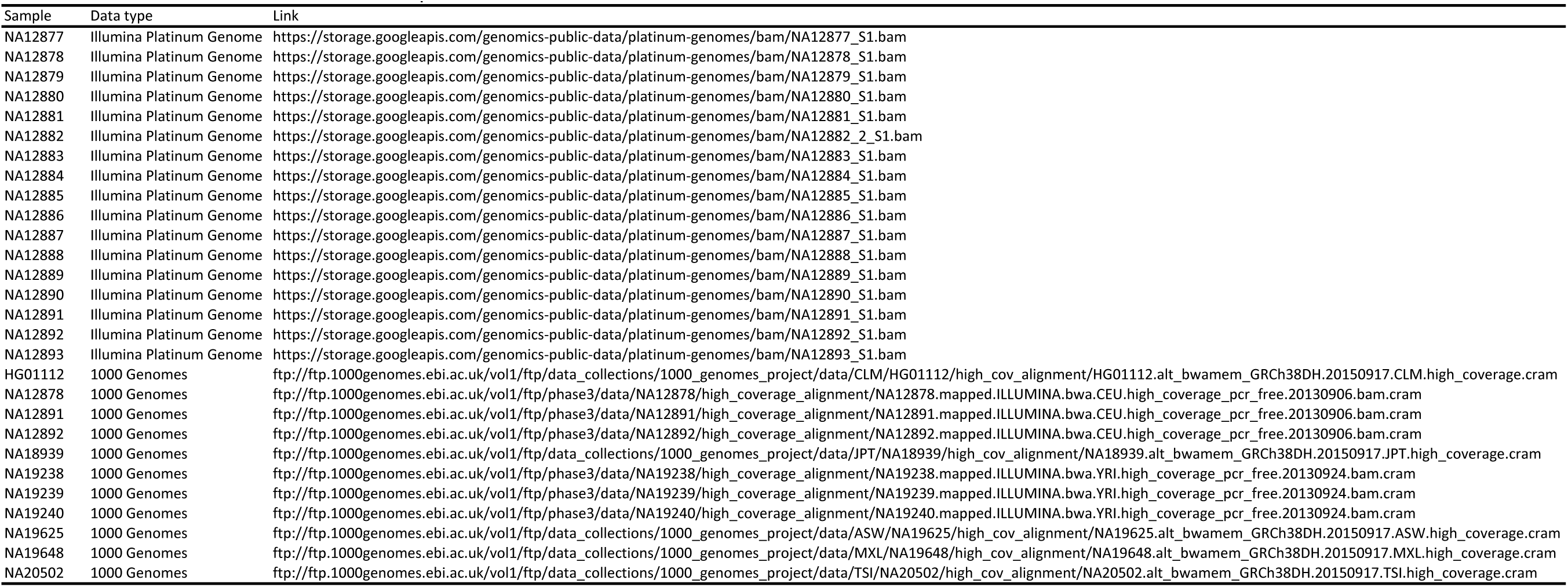
List of Illumina Platinum Genome and 1000 Genomes samples used and their URLs.

**S3.**
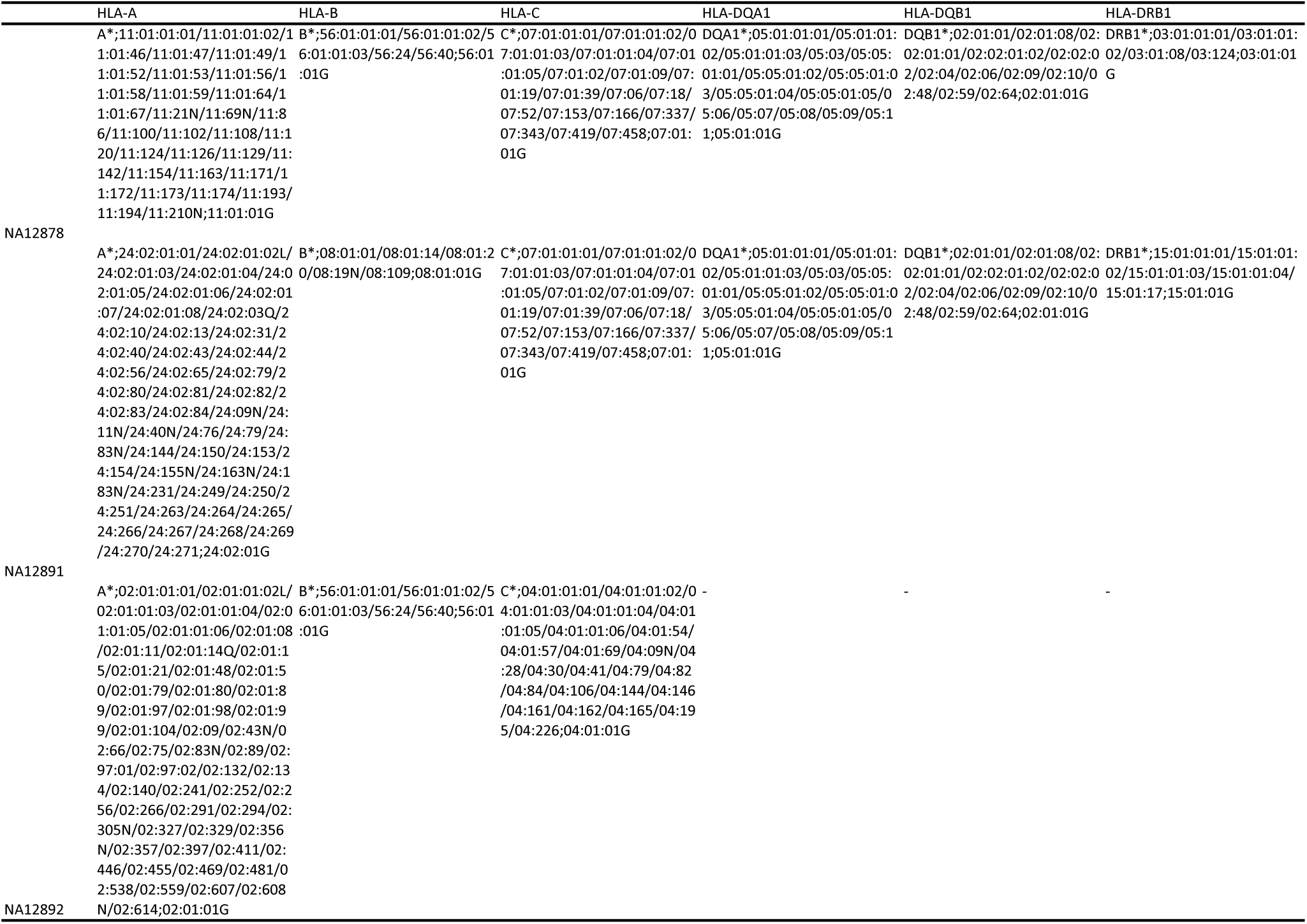
List of removed alleles and its ‘G’ group allele name from Platinum Trio (NA12878-NA12891-NA1892). Each row represents a individual in the trio and the columns are 6 HLA genes typed by Kourami.

**S4.**
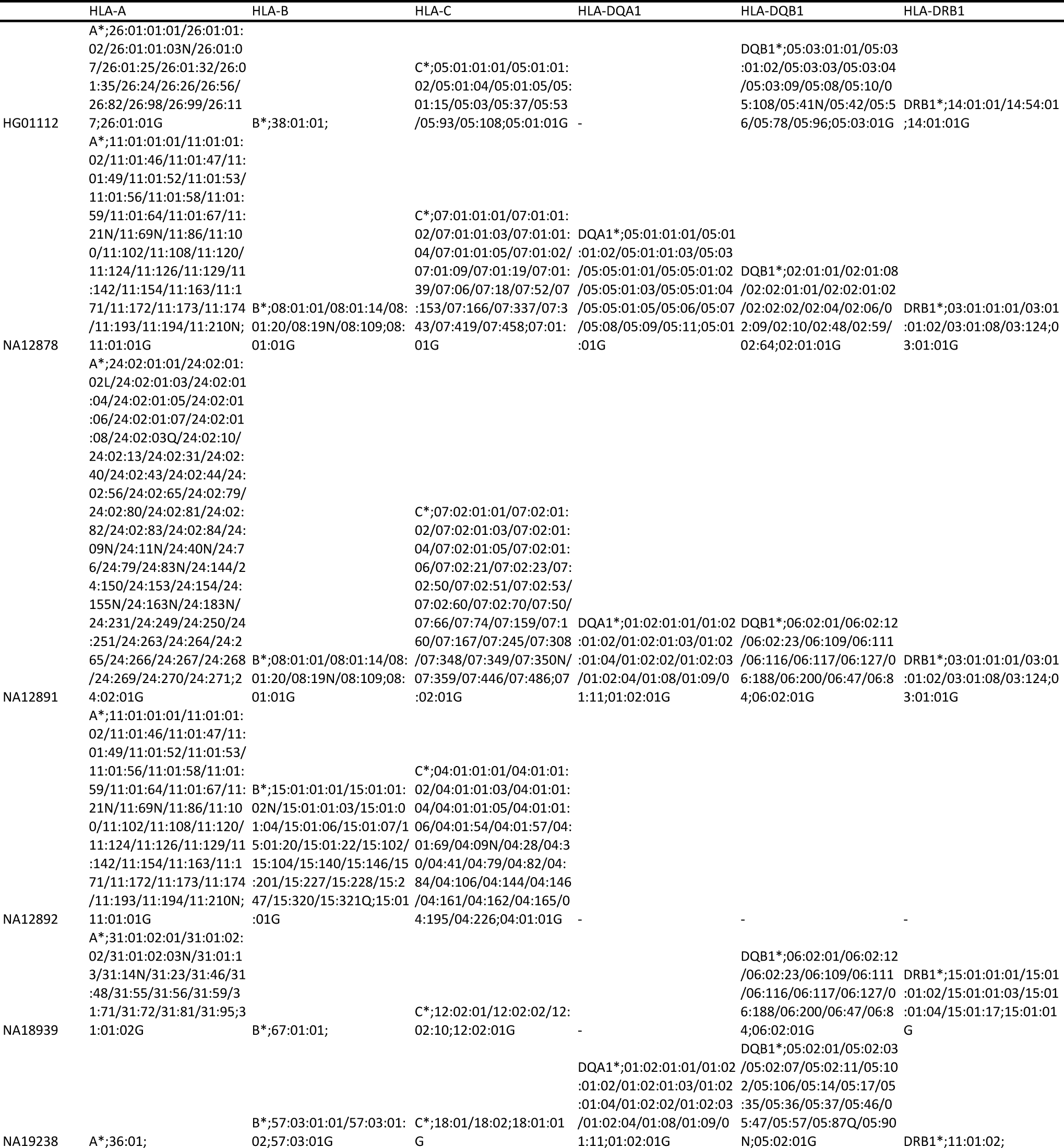
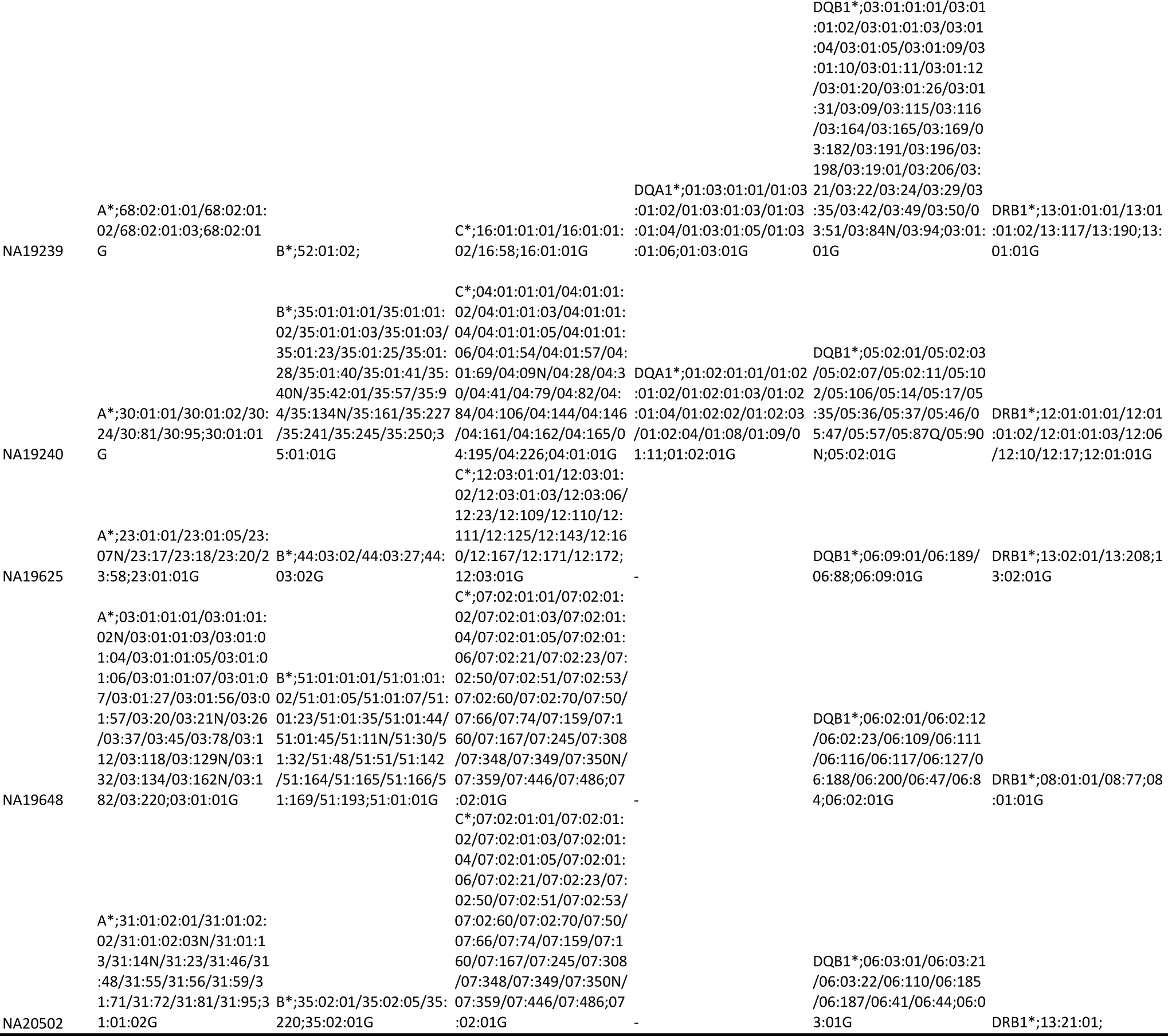
List of removed alleles and its ‘G’ group allele name from 1000 Genomes data. Each row represents an individual from 1000 Genome Project and the columns are 6 HLA genes typed by Kourami.

**S5.** List of removed ‘G’ group alleles from Simulation data (100 replicates). Each row represents a simulation replicate and the columns represents the 6 HLA genes typed by Kourami. Missing ‘-’ allele is present if no allele can be removed because the correct allele for the replicate is the reference allele for the given multiple sequence alignment in the databse.

| Simulation Experiment | HLA-A | HLA-B | HLA-C | HLA-DQA1 | HLA-DQB1 | HLA-DRB1 |
| --- | --- | --- | --- | --- | --- | --- |
| Simul0 | A*02:03:01G | B*15:142 | C*02:16:02 | DQA1*04:01:01G | DQB1*03:03:02G | DRB1*07:01:01G |
| Simul1 | A*30:02:01G | B*15:42 | C*06:02:01G | DQA1*06:01:01G | DQB1*03:03:02G | DRB1*11:01:02 |
| Simul2 | A*02:01:01G | B*51:07:01 | C*17:01:01G | DQA1*03:01:01G | DQB1*03:05:01G | DRB1*12:01:01G |
| Simul3 | A*24:02:01G | B*35:01:01G | C*07:02:64 | DQA1*02:01:01G | DQB1*06:03:20 | DRB1*12:01:01G |
| Simul4 | A*34:02:01 | B*40:309 | C*07:02:64 | DQA1*04:01:01G | DQB1*06:09:01G | DRB1*09:01:02G |
| Simul5 | A*01:11N | B*40:309 | C*06:149 | DQA1*04:01:01G | DQB1*03:05:01G | DRB1*11:04:01G |
| Simul6 | A*74:01:01G | B*37:52 | C*01:99 | DQA1*01:02:01G | DQB1*06:09:01G | DRB1*15:01:01G |
| Simul7 | A*68:01:01G | B*40:130:02 | C*08:21 | DQA1*04:01:01G | DQB1*02:62 | DRB1*08:03:02 |
| Simul8 | A*03:36N | B*38:01:01 | C*08:73 | DQA1*01:03:01G | DQB1*03:01:01G | DRB1*15:01:01G |
| Simul9 | A*24:07:01 | B*38:11 | C*01:106 | DQA1*01:02:01G | DQB1*02:01:01G | DRB1*16:02:01G |
| Simul10 | A*02:32N | B*08:08N | C*02:02:02G | DQA1*06:01:01G | DQB1*02:01:01G | DRB1*15:02:01G |
| Simul11 | A*74:01:03 | B*37:01:05 | C*03:02:01G | DQA1*05:01:01G | DQB1*03:01:01G | DRB1*01:02:01 |
| Simul12 | A*02:193 | B*56:01:01G | C*02:10:01G | - | DQB1*06:190 | DRB1*15:01:01G |
| Simul13 | A*02:03:01G | B*07:05:01G | C*04:01:01G | DQA1*04:01:01G | DQB1*03:03:02G | DRB1*01:02:01 |
| Simul14 | A*03:01:01G | B*07:252 | C*07:149 | DQA1*01:02:01G | DQB1*06:03:01G | DRB1*15:03:01G |
| Simul15 | A*11:02:01G | B*35:01:01G | C*03:192 | DQA1*03:01:01G | DQB1*03:03:02G | DRB1*11:01:01G |
| Simul16 | A*24:02:01G | B*39:104 | C*08:12 | DQA1*01:02:01G | DQB1*03:03:02G | DRB1*01:02:01 |
| Simul17 | A*31:01:02G | B*07:05:01G | C*07:02:64 | DQA1*01:02:01G | DQB1*03:03:02G | DRB1*03:01:01G |
| Simul18 | A*33:07 | B*27:144 | C*14:69 | DQA1*05:01:01G | DQB1*05:03:01G | DRB1*15:02:01G |
| Simul19 | A*26:01:01G | B*15:02:01G | C*08:21 | DQA1*01:02:01G | DQB1*02:53Q | DRB1*15:01:01G |
| Simul20 | A*24:07:01 | B*18:02 | C*17:01:01G | DQA1*05:01:01G | DQB1*06:02:01G | DRB1*14:01:01G |
| Simul21 | A*31:01:02G | B*55:02:01G | C*07:01:45 | DQA1*06:01:01G | DQB1*02:53Q | DRB1*15:01:01G |
| Simul22 | A*01:09 | B*40:02:01G | C*12:155Q | DQA1*01:02:01G | DQB1*06:02:25 | DRB1*11:01:01G |
| Simul23 | A*02:376 | B*39:64 | C*04:15:02 | DQA1*01:10 | DQB1*03:03:02G | DRB1*15:01:01G |
| Simul24 | A*01:02 | B*08:144 | C*03:40:01G | DQA1*05:01:01G | DQB1*06:02:25 | DRB1*15:01:01G |
| Simul25 | A*68:01:02G | B*08:26:03 | C*08:113 | DQA1*01:02:01G | DQB1*06:03:20 | DRB1*11:01:02 |
| Simul26 | A*02:269 | B*13:15 | C*02:102 | DQA1*05:01:01G | DQB1*02:62 | DRB1*14:05:01 |
| Simul27 | A*02:51 | B*14:01:01 | C*07:422 | DQA1*04:01:01G | DQB1*02:01:01G | DRB1*15:03:01G |
| Simul28 | A*26:50 | B*44:218 | C*07:392 | DQA1*01:02:01G | DQB1*02:01:01G | DRB1*11:01:01G |
| Simul29 | A*02:07:01G | B*78:01:01 | C*01:99 | DQA1*01:10 | DQB1*06:01:01G | DRB1*15:02:01G |
| Simul30 | A*32:06 | B*35:05:01 | C*03:02:01G | DQA1*01:10 | DQB1*06:03:01G | DRB1*10:01:01G |
| Simul31 | A*24:61 | B*13:02:01G | C*08:01:01G | DQA1*01:02:01G | DQB1*06:125 | DRB1*01:02:01 |
| Simul32 | A*29:01:01G | B*39:64 | C*07:01:45 | DQA1*03:01:01G | DQB1*06:02:01G | DRB1*12:01:01G |
| Simul33 | A*24:152 | B*38:01:01 | C*18:01:01G | DQA1*05:01:01G | DQB1*03:03:02G | DRB1*15:03:01G |
| Simul34 | A*32:69 | B*44:64:02 | C*01:99 | DQA1*01:02:01G | DQB1*06:03:01G | DRB1*15:01:01G |
| Simul35 | A*66:01:01G | B*51:01:02 | C*06:153 | DQA1*01:02:01G | DQB1*05:02:01G | DRB1*08:03:02 |
| Simul36 | A*03:02:01G | B*08:56:02 | C*15:17 | DQA1*06:01:01G | DQB1*03:02:01G | DRB1*13:02:01G |
| Simul37 | A*68:02:01G | B*35:271 | C*03:02:01G | DQA1*03:01:01G | DQB1*03:03:02G | DRB1*10:01:01G |
| Simul38 | A*11:183 | B*53:01:01G | C*05:113N | DQA1*04:01:01G | DQB1*06:190 | DRB1*12:01:01G |
| Simul39 | A*68:07 | B*55:01:01G | C*15:05:01G | DQA1*02:01:01G | DQB1*05:03:01G | DRB1*09:01:02G |
| Simul40 | A*01:03 | B*35:05:01 | C*06:153 | DQA1*01:07Q | DQB1*06:09:01G | DRB1*15:02:01G |
| Simul41 | A*29:02:01G | B*51:01:02 | C*06:02:01G | DQA1*05:01:01G | DQB1*06:03:01G | DRB1*10:01:01G |
| Simul42 | A*02:01:01G | B*46:01:01G | C*04:187 | DQA1*06:01:01G | DQB1*03:02:01G | DRB1*11:01:01G |
| Simul43 | A*74:01:01G | B*52:31:01 | C*03:290 | DQA1*02:01:01G | DQB1*06:03:20 | DRB1*07:01:01G |
| Simul44 | A*34:01:01 | B*18:69 | C*03:06:01 | DQA1*03:01:01G | DQB1*03:03:02G | DRB1*16:02:01G |
| Simul45 | A*11:01:01G | B*39:92 | C*07:32N | DQA1*01:02:01G | DQB1*03:02:01G | DRB1*11:01:01G |
| Simul46 | A*02:60:01 | B*56:01:01G | C*12:03:34G | DQA1*01:03:01G | DQB1*02:62 | DRB1*12:01:01G |
| Simul47 | A*24:03:01G | B*07:253 | C*14:21N | DQA1*01:03:01G | DQB1*06:02:01G | DRB1*07:01:01G |
| Simul48 | A*02:06:01G | B*35:95 | C*07:19 | DQA1*03:01:01G | DQB1*06:02:25 | DRB1*09:01:02G |
| Simul49 | A*31:01:04 | B*39:13:02 | C*17:17 | DQA1*03:01:01G | DQB1*06:190 | DRB1*15:01:01G |
| Simul50 | A*11:74 | B*18:01:01G | C*03:250 | DQA1*05:01:01G | DQB1*03:03:02G | DRB1*15:01:01G |
| Simul51 | A*24:07:01 | B*35:01:22 | C*07:02:64 | DQA1*01:02:01G | DQB1*06:03:20 | DRB1*12:01:01G |
| Simul52 | A*24:02:01G | B*38:14 | C*07:56:02 | DQA1*05:01:01G | DQB1*03:01:01G | DRB1*08:03:02 |
| Simul53 | A*02:02:01G | B*56:01:01G | C*06:156 | DQA1*06:01:01G | DQB1*03:05:01G | DRB1*11:04:01G |
| Simul54 | A*01:194 | B*07:255 | C*02:02:02G | DQA1*01:02:01G | DQB1*06:02:25 | DRB1*10:01:01G |
| Simul55 | A*33:07 | B*44:225 | C*02:02:02G | DQA1*01:10 | DQB1*06:03:01G | DRB1*14:05:01 |
| Simul56 | A*24:02:01G | B*35:300 | C*08:103 | - | DQB1*02:62 | DRB1*07:01:01G |
| Simul57 | A*24:02:01G | B*08:20 | C*03:04:48 | DQA1*01:10 | DQB1*05:03:01G | DRB1*07:01:01G |
| Simul58 | A*31:01:02G | B*15:32:01 | C*04:128 | DQA1*05:01:01G | DQB1*03:02:01G | DRB1*11:01:02 |
| Simul59 | A*02:06:01G | B*35:41 | C*07:386 | DQA1*05:01:01G | DQB1*03:01:01G | DRB1*10:01:01G |
| Simul60 | A*01:02 | B*40:305 | C*04:01:01G | DQA1*05:01:01G | DQB1*03:05:01G | DRB1*15:02:01G |
| Simul61 | A*29:01:01G | B*35:08:01 | C*07:60 | DQA1*05:01:01G | DQB1*05:02:01G | DRB1*14:01:01G |
| Simul62 | A*02:533 | B*58:01:01G | C*02:86 | DQA1*05:01:01G | DQB1*02:01:01G | DRB1*09:01:02G |
| Simul63 | A*02:60:01 | B*57:05 | C*07:26:01 | - | DQB1*03:02:01G | DRB1*15:01:01G |
| Simul64 | A*03:01:01G | B*27:25 | C*12:99 | DQA1*01:02:01G | DQB1*06:03:20 | DRB1*14:05:01 |
| Simul65 | A*03:213 | B*08:144 | C*12:03:37 | DQA1*04:01:01G | DQB1*03:01:01G | DRB1*07:01:01G |
| Simul66 | A*68:113 | B*40:10:01G | C*02:92N | DQA1*01:07Q | DQB1*06:09:01G | DRB1*15:01:01G |
| Simul67 | A*68:71 | B*27:24 | C*07:01:01G | DQA1*01:02:01G | DQB1*06:01:01G | DRB1*13:01:01G |
| Simul68 | A*31:01:23 | B*44:220 | C*01:99 | DQA1*01:03:01G | DQB1*03:05:01G | DRB1*12:01:01G |
| Simul69 | A*29:02:17 | B*27:05:02G | C*01:40 | DQA1*05:01:01G | DQB1*03:03:02G | DRB1*11:01:02 |
| Simul70 | A*24:02:01G | B*15:25:01G | C*01:106 | DQA1*01:03:01G | DQB1*02:01:01G | DRB1*11:01:02 |
| Simul71 | A*02:01:01G | B*15:25:01G | C*05:110 | DQA1*03:01:01G | DQB1*03:01:01G | DRB1*15:03:01G |
| Simul72 | A*24:02:01G | B*73:01 | C*04:01:01G | DQA1*02:01:01G | DQB1*02:01:01G | DRB1*11:01:01G |
| Simul73 | A*31:01:02G | B*14:07N | C*03:287 | DQA1*04:01:01G | DQB1*02:62 | DRB1*08:03:02 |
| Simul74 | A*29:03 | B*38:01:08 | C*04:01:01G | DQA1*04:01:01G | DQB1*02:53Q | DRB1*01:02:01 |
| Simul75 | A*29:02:17 | B*41:37 | C*02:85 | DQA1*03:01:01G | DQB1*05:03:01G | DRB1*03:01:01G |
| Simul76 | A*74:01:01G | B*67:01:01 | C*07:205 | DQA1*03:01:01G | DQB1*05:03:01G | DRB1*15:03:01G |
| Simul77 | A*24:07:01 | B*27:05:18 | C*04:01:01G | DQA1*04:01:01G | - | DRB1*01:02:01 |
| Simul78 | A*23:01:01G | B*49:01:01G | C*08:90 | DQA1*01:02:01G | DQB1*02:53Q | DRB1*15:01:01G |
| Simul79 | A*68:17 | B*47:01:01G | C*03:02:01G | DQA1*01:02:01G | DQB1*03:03:02G | DRB1*11:04:01G |
| Simul80 | A*24:252N | B*44:02:01G | C*17:30 | DQA1*05:01:01G | DQB1*02:01:01G | DRB1*03:01:01G |
| Simul81 | A*24:07:01 | B*40:10:01G | C*06:160 | DQA1*03:01:01G | DQB1*03:03:02G | DRB1*14:05:01 |
| Simul82 | A*01:20 | B*55:76 | C*02:103 | DQA1*05:01:01G | DQB1*03:03:02G | DRB1*03:01:01G |
| Simul83 | A*26:11N | B*35:285 | C*03:04:01G | DQA1*01:02:01G | DQB1*03:03:02G | DRB1*10:01:01G |
| Simul84 | A*68:71 | B*39:92 | C*12:19 | DQA1*03:01:01G | DQB1*06:125 | DRB1*12:01:01G |
| Simul85 | A*24:03:01G | B*48:04:01 | C*07:30 | DQA1*05:01:01G | DQB1*03:01:01G | DRB1*15:02:01G |
| Simul86 | A*02:05:01G | B*39:01:01G | C*05:08 | DQA1*04:01:01G | DQB1*06:125 | DRB1*15:02:01G |
| Simul87 | A*01:11N | B*51:01:02 | C*07:412 | DQA1*04:01:01G | DQB1*03:01:01G | DRB1*15:01:01G |
| Simul88 | A*02:01:01G | B*55:12 | C*06:148 | DQA1*03:01:01G | DQB1*06:09:01G | DRB1*04:03:01 |
| Simul89 | A*68:08:01 | B*07:02:48 | C*04:71 | DQA1*01:02:01G | DQB1*02:62 | DRB1*07:01:01G |
| Simul90 | A*74:01:01G | B*35:05:01 | C*12:03:01G | DQA1*04:01:01G | DQB1*03:05:01G | DRB1*13:02:01G |
| Simul91 | A*02:06:01G | B*15:01:38 | C*02:81 | DQA1*01:02:01G | DQB1*06:03:20 | DRB1*15:01:01G |
| Simul92 | A*31:01:04 | B*13:02:17 | C*14:02:19 | DQA1*06:01:01G | DQB1*06:02:01G | DRB1*15:02:01G |
| Simul93 | A*80:01:01G | B*55:48 | C*05:08 | DQA1*01:02:01G | DQB1*02:01:01G | DRB1*12:01:01G |
| Simul94 | A*68:07 | B*08:01:01G | C*15:02:01G | DQA1*05:01:01G | DQB1*03:05:01G | DRB1*12:01:01G |
| Simul95 | A*74:01:01G | B*51:01:01G | C*17:01:01G | DQA1*03:01:01G | DQB1*03:02:01G | DRB1*14:01:01G |
| Simul96 | A*24:03:01G | B*82:02:01 | C*04:201 | DQA1*05:01:01G | DQB1*03:02:01G | DRB1*15:01:01G |
| Simul97 | A*26:01:39 | B*42:02:01G | C*03:04:04 | DQA1*01:03:01G | DQB1*02:01:01G | DRB1*11:04:01G |
| Simul98 | A*24:02:01G | B*15:123:01G | C*01:02:29 | DQA1*05:01:01G | DQB1*06:03:01G | DRB1*07:01:01G |
| Simul99 | A*32:69 | B*40:03:01G | C*07:01:01G | DQA1*03:01:01G | DQB1*03:01:01G | DRB1*16:02:01G |

**S6.** Kourami’s typing accuracy with varing sequencing coverages. A total of 5 replicates of random sampling at each varying coverage was used. 2 alleles per each replicate per each of 6 HLA genes make up a total of 60 calls.

| Coverage | Number of correct calls | Total number of calls | Accuracy |
| --- | --- | --- | --- |
| 40x | 60 | 60 | 1.00 |
| 30x | 58 | 60 | 0.97 |
| 20x | 49 | 60 | 0.82 |

